# YAP and TAZ regulate adherens junction dynamics and endothelial cell distribution during vascular development

**DOI:** 10.1101/174185

**Authors:** Filipa Neto, Alexandra Klaus-Bergmann, Yu-Ting Ong, Silvanus Alt, Anne-Clémence Vion, Anna Szymborska, Joana R. Carvalho, Irene Hollfinger, Eireen Bartels-Klein, Claudio A. Franco, Michael Potente, Holger Gerhardt

## Abstract

Formation of a hierarchically organized blood vessel network by sprouting angiogenesis is critical for tissue growth, homeostasis and regeneration. How in this process endothelial cells arise in adequate numbers and arrange suitably to shape a functional vascular network is poorly understood. Here we show that YAP and TAZ promote stretch-induced proliferation and rearrangements of endothelial cells whilst preventing bleeding in developing vessels. Mechanistically, YAP and TAZ increase VE-cadherin turnover at junctions and suppress endothelial Notch and BMP signaling, two key pathways that limit sprouting and endothelial dynamics. Consequently, the loss of YAP and TAZ leads to stunted sprouting with local aggregation as well as scarcity of endothelial cells, branching irregularities and junction defects. Forced nuclear activity of TAZ instead drives hypersprouting and vascular hyperplasia. We propose a new model in which YAP and TAZ integrate mechanical signals with Notch and BMP signaling to balance endothelial cell distribution in angiogenic vessels.

## INTRODUCTION

A long-standing question in developmental and cell biology relates to how cells integrate mechanical and chemical signals to orchestrate the morphogenic behaviours that ensure adequate tissue patterning. During sprouting angiogenesis, the arrangement and distribution of cells rather than their numbers appear to drive morphogenesis of the vascular tree. Recent data showing unaltered remodelling in the absence of endothelial cell apoptosis and normal branching frequency across a range of endothelial cell densities support this idea (1). In the extreme, however, too few cells will jeopardize network formation and stability (2), whereas too many cells might compromise vessel calibre control (1). Functional network formation therefore needs to establish the right number of cells in the right place, and distribute them such that the hierarchical branching pattern is supported. What establishes such a balance has remained unclear. Here we provide evidence for the yes-associated protein 1 (YAP) and its paralog WW domain containing transcription regulator 1 (TAZ) to be critically involved as endothelial cell autonomous regulators in this process.

YAP and TAZ, two transcriptional co-activators initially discovered as effectors of the Hippo signalling pathway, play a central role in organ size control via regulation of proliferation and apoptosis (3-5). In confluent cells, YAP and TAZ are phosphorylated by the Hippo kinase cascade, and retained in the cytoplasm. In sparse cells, YAP and TAZ remain unphosphorylated and can translocate to the nucleus, where they bind transcription factors inducing the expression of pro-proliferative and anti-apoptotic genes. Other stimuli have been found to regulate YAP and TAZ nuclear translocation and activity – these include, among others, G-protein coupled receptors (GPCRs) (6), junctional proteins (7, 8), and mechanical stimuli (9, 10). Furthermore, besides cell proliferation and apoptosis, YAP and TAZ also regulate cell differentiation (11), migration (12) and actomyosin contraction (13). In vascular development, the roles of YAP and TAZ are not fully understood. *Yap* null mutant zebrafish develop an initially normal vasculature but display increased vessel collapse and regression. *Yap/Taz* double mutant zebrafish die before the onset of circulation with severe developmental defects, precluding analysis of vascular development in this context (14). Endothelial-specific deletion of *Yap* in mice using the Tie2-Cre transgenic line is embryonically lethal due to heart valve defects caused by failed endothelial-to-mesenchymal transition (15). During post-natal development of the mouse retina, YAP was shown to regulate vascular density and branching by promoting the transcription of *Angiopoetin-2* (16). While these studies point towards an important role for YAP in regulating blood vessel formation and maintenance, none of them addressed the endothelial cell autonomous requirement for YAP during sprouting angiogenesis. In addition, no study has addressed a potential requirement for TAZ during angiogenesis, and the possible redundancy between both proteins in this context. Finally, the possible interplay between YAP/TAZ and the major signalling pathways regulating angiogenesis has not been assessed.

Here, we show that YAP and TAZ are both expressed and active in sprouting ECs and critical for sprouting angiogenesis. The inducible, endothelial-specific combined deletion of YAP and TAZ leads to severe morphogenic defects consistent with impaired junctional remodelling *in vivo.* Furthermore, we found that the loss of YAP and TAZ decreased VE-Cadherin turnover and increased cell-cell coupling. We also discovered that endothelial YAP and TAZ strongly regulate endothelial Notch and BMP signalling *in vitro* and *in vivo,* together suggesting that YAP and TAZ integrate mechanical stimuli with key transcriptional regulators of endothelial sprouting and cell rearrangements during angiogenesis.

## RESULTS

### YAP and TAZ have distinct expression patterns in endothelial cells of developing vessels and localise to the nucleus at the sprouting front

Immunofluorescence staining in the postnatal mouse retina showed that YAP and TAZ are distinctly expressed in the ECs of the developing vasculature (Figure 1). While YAP is evenly expressed throughout the vasculature (Figure 1 A-D), the expression of TAZ is especially prominent at the sprouting front (Figure 1 – E-H). Furthermore, YAP is exclusively cytoplasmic in all areas of the retinal vasculature, with the exception of the sprouting front where some ECs express nuclear YAP, although at lower levels than in the cytoplasm. (Figure 1A’-D’). TAZ staining signal is very low in the remodelling plexus, arteries and veins (Figure1 E’-H’); at the sprouting front, TAZ is strongly nuclear in numerous ECs (Figure 1 E, green arrowheads and E’), and both nuclear and cytoplasmic in others (Figure 1E, red arrowheads). The nuclear signal of YAP and TAZ did not correlate with a tip or stalk cell phenotype; nuclear YAP and TAZ are rather present in a subset of tip and stalk ECs at the sprouting front. YAP and TAZ were also found at endothelial adherens junctions in veins and in the remodelling plexus, (yellow arrowheads in Fig. 1D’ and F’), as revealed by co-staining for VE-Cadherin (Figure 1 – Figure supplement 1). Together, these observations suggest that YAP/TAZ are abundant transcriptional co-activators in the endothelium, and dynamically regulated during the angiogenic process.

**Figure 1.**
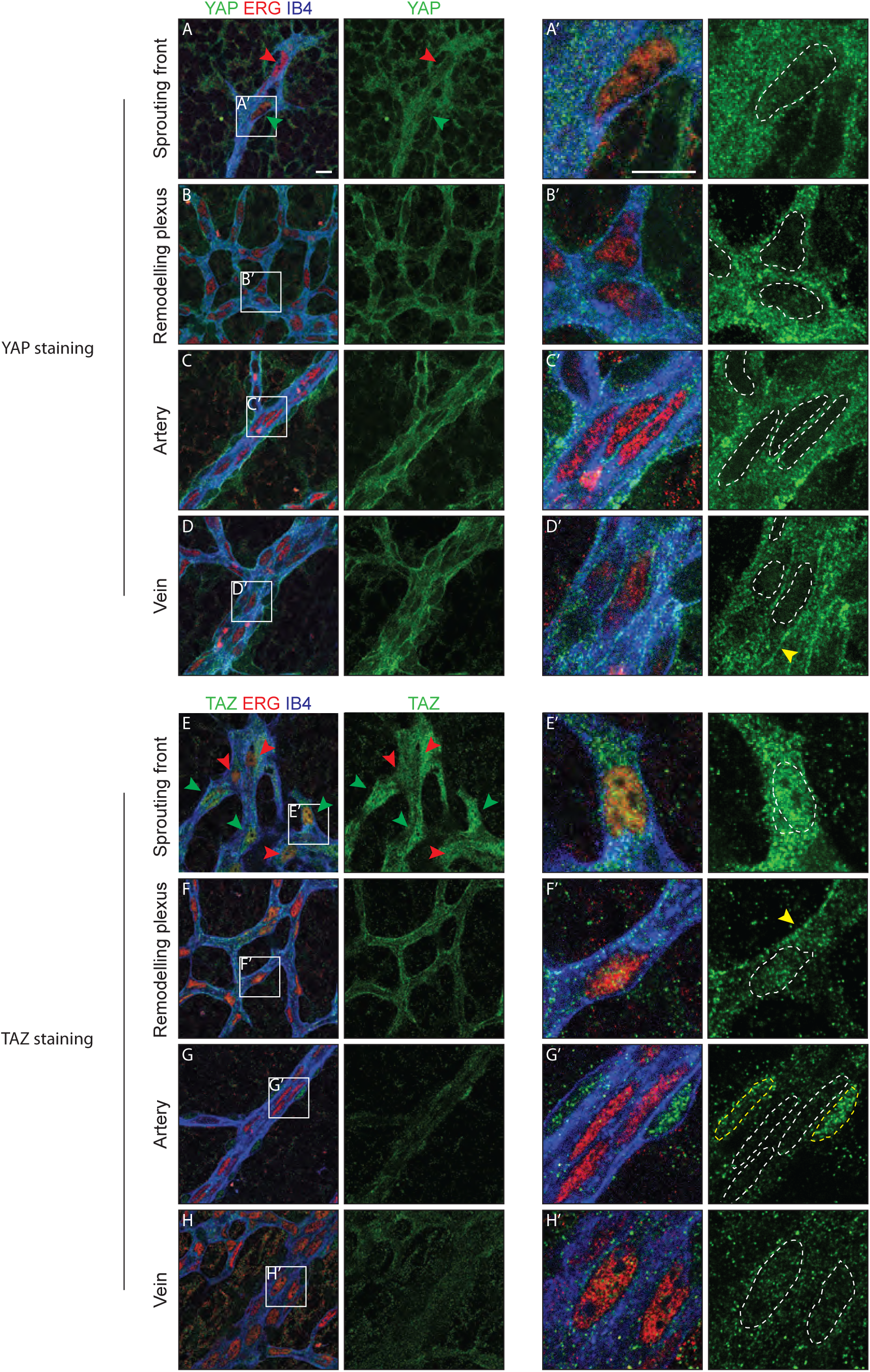
YAP and TAZ are expressed throughout the vasculature of developing mouse retinas, and localise to the nucleus of sprouting endothelial cells. Immunofluorescence staining of YAP (green, A-D and A’-D’) and TAZ (green, E-H and E’-H’) was performed in wild-type mouse retinas at post-natal day 6 (P6). Retinas were co-stained with the endothelial membrane marker Isolectin-B4 (IB4; blue) and with antibodies against the endothelial nuclei marker ERG (red). White dotted lines, outline of endothelial nuclei. Yellow dotted lines, outline of perivascular cells’ nuclei. Green arrowheads, nuclear localisation of YAP and TAZ. Red arrowheads, cytoplasmic localisation of YAP and TAZ. Yellow arrowheads, junctional localisation of YAP and TAZ. Images correspond to single confocal planes. Scale bar: 10μm.

### YAP and TAZ are required for vessel growth, branching and regularity of the vasculature

To examine the cell-autonomous role of endothelial YAP and TAZ during angiogenesis we crossed mice bearing *floxed* alleles of *Yap* or *Taz* (17) with mice expressing a tamoxifen-inducible Cre recombinase driven by the endothelial-restricted *Pdgfb* promoter (*Pdgfb-iCreERT2*) (18). Injection of the offspring with tamoxifen induced loss of YAP and TAZ protein in ECs during post-natal vascular development, as evidenced by immunofluorescence staining (Figure 2 – Figure supplement 1). Cre negative littermate mice were used as controls.

Endothelial deletion of YAP or TAZ led to mild vascular defects (Figure 2A,B,C,D). *Yap*^fl/fl^ *Pdgfb-iCreERT2* mice (*Yap* iEC-KO) presented reduced radial expansion of the vasculature (7% +/- 5.4 reduction, *p*=0.0123) and reduced vessel density (9% +/- 4.4 reduction, *p*=0.0002) (Figure 2G,H). *Taz*^fl/fl^ *Pdgfb-iCreERT2* mice (*Taz* iEC-KO) did not show altered radial expansion but displayed decreased vessel density (6% +/- 5.8 reduction, *p*=0.0214) (Figure 2H). Neither mutant showed a change in the branching frequency of vessels (Figure 2I). Interestingly, in *Yap* iEC-KO retinas the expression of TAZ was increased and TAZ more often localised to the nucleus (Figure 2 – Figure supplement 2), suggesting compensatory regulation. *Taz* iEC-KO retinas did not however show a clear difference in YAP expression (data not shown). Deleting both proteins in compound mutant mice (*Yap*^fl/f^ *Taz*^fl/fl^ *Pdgfb-iCreERT2*, *YapTaz* iEC-KO) produced a dramatic defect in blood vessel development (Figure 2E,F): the retinal vasculature showed a 21% (+/-14, *p*=0.0012) decrease in radial expansion (Figure 2G), a 26% (+/- 7.0, *p*<0.0001) decrease in capillary density (Figure 2H), and a 55% (+/- 15.4, *p*<0.0001) decrease in branching frequency (Figure 2I). Interestingly, the vessel loops were not only bigger in *Yap/Taz* iEC-KO mice (Figure 2J), but also more variable in size (Figure 2K), and shape (Figure 2L) than in control mice. These results indicate that endothelial YAP and TAZ are critical for the development of a homogeneous blood vessel network and can perform redundant functions in the endothelium.

**Figure 2.**
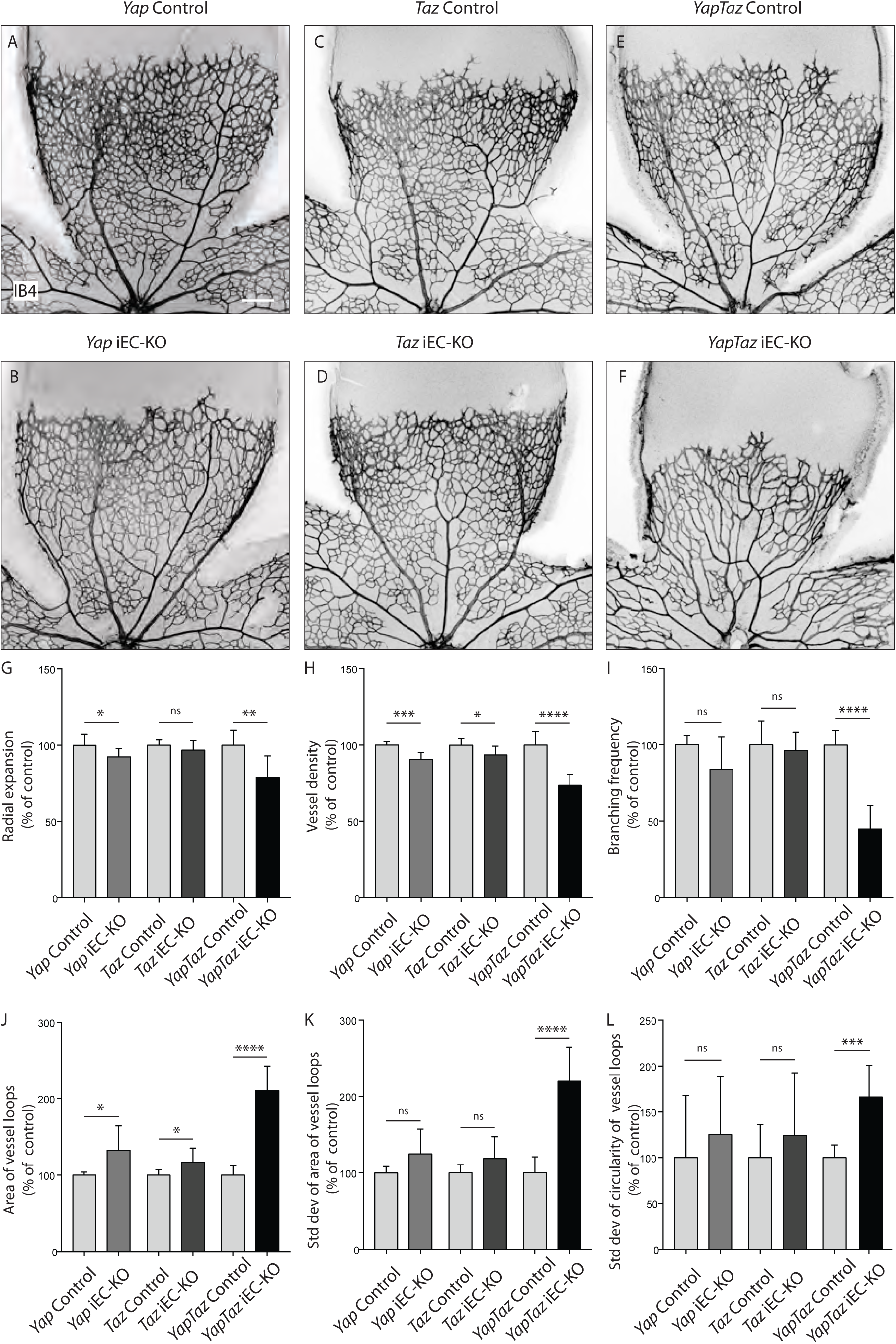
Endothelial YAP and TAZ are required for vessel growth, branching and homogeneity of the plexus. (**A-F**) Retinas from P6 *Yap* iEC-KO (B), *Taz* iEC-KO (D) and *YapTaz* iEC-KO (F), and respective control pups (A,C,E) were stained with Isolectin B4 (IB4). Scale bar: 200μm. (**G-J**) Quantification of radial expansion (G), vessel density (H), branching frequency (I) and area of vessel loops (J) in *Yap* iEC-KO, *Taz* iEC-KO and *YapTaz* iEC-KO. Results are shown as percentage of the respective controls. Data are mean +/- SD. n≥ 5 pups. *p* values were calculated using unpaired t-test. *, *p*<0.05; **, *p*<0.01; ****, *p*<0.0001. (**K-L**) Quantification of the standard deviation of the area (K) and circularity (L) of the vessels loops in *Yap* iEC-KO, *Taz* iEC-KO and *YapTaz* iEC-KO retinas. Results are shown as percentage of the respective controls. Data are mean +/- SD. n≥ 5 pups. *p* values were calculated using unpaired t-test. *, *p*<0.05; **, *p*<0.01; ***, *p*<0.001****, *p*<0.0001.

### YAP and TAZ are required for endothelial cell proliferation in response to mechanical stretch

As YAP and TAZ display pro-proliferative and anti-apoptotic roles in many cell types (3, 4), we evaluated whether the reduced vascularization of *Yap/Taz* iEC-KO retinas was associated with reduced cell proliferation or increased apoptosis. EC proliferation, assessed by EdU staining (Figure 3A-C), was decreased in *Yap* iEC-KO retinas (23% +/- 10.0, *p*=0.0469), whilst not affected in *Taz* iEC-KO. Consistent with our prior results the decrease in cell proliferation was strongest in *Yap/Taz* iEC-KO retinas (33% +/- 26.0, *p*=0.0059). Staining for cleaved caspase 3 revealed that apoptosis was unaltered by YAP/TAZ loss (Figure 3D-F).

**Figure 3.**
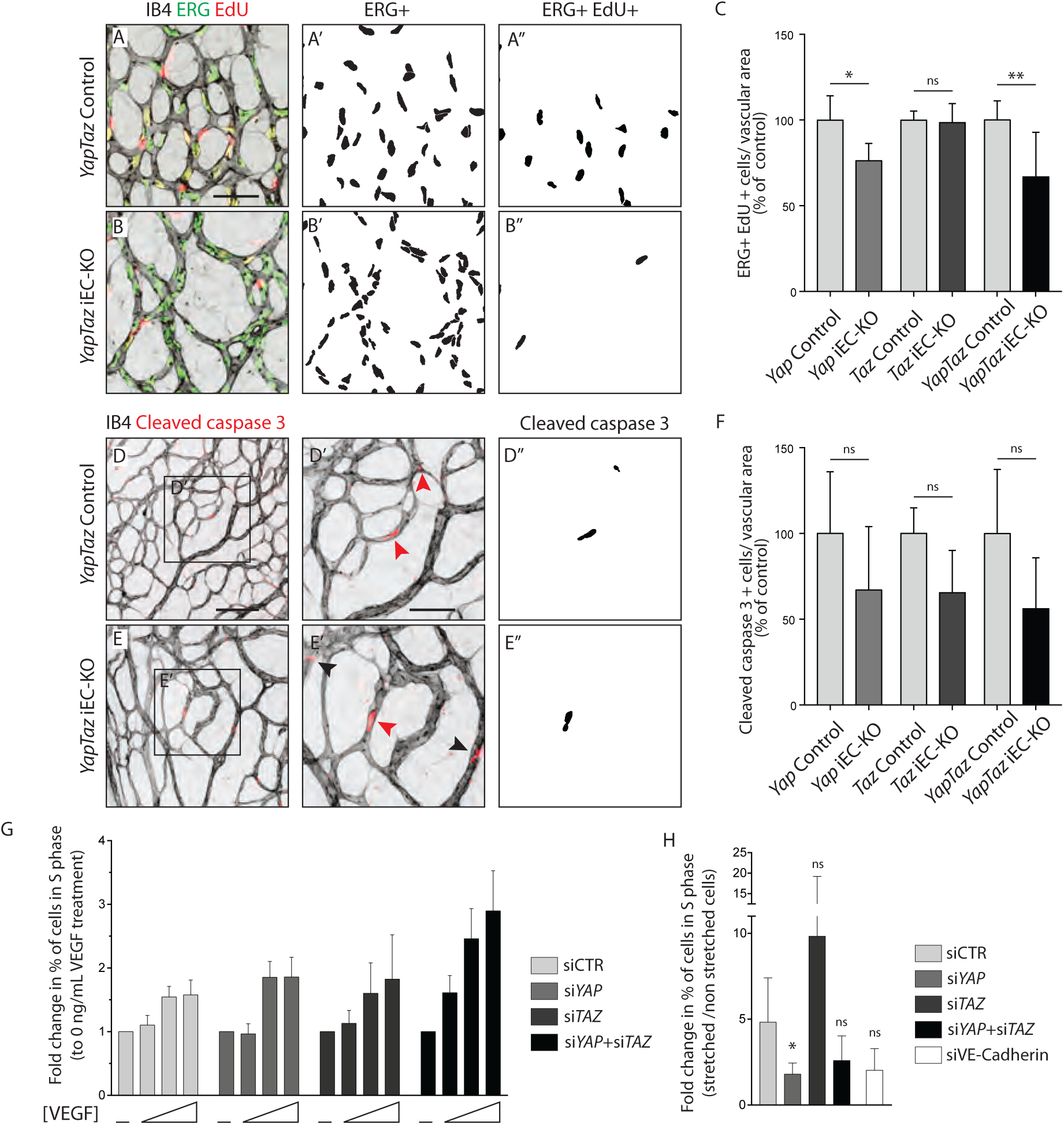
YAP and TAZ are required for endothelial cell proliferation *in vivo* and endothelial cell proliferation in response to mechanical stretch *in vitro*. **A,B**, P6 retinal vessels labelled with IB4 (grey) and stained for EdU (red, marking S phase positive cells) and Erg (green, marking endothelial nuclei) in *YapTaz* iEC-KO (**B**) and littermate control mice (**A**). **A’**,**B’**, mask of Erg + cells indicating endothelial nuclei. **A”**, **B”**, mask of Erg + and EdU + cells indicating proliferating endothelial cells. **C**, Quantification of endothelial proliferation in *Yap* iEC-KO (n=3 control/4 KO pups), *Taz* iEC-KO (n=5 control/5 KO pups) and *YapTaz* iEC-KO (n=8 control/7 KO pups). Number of EdU-positive and ERG-positive cells per IB4 labelled vascular area was calculated for each genotype and results are shown in percentage of the respective controls. Data are mean +/- SD. *p* values were calculated using unpaired *t*-test. ns, *p*>0.05; *, *p*<0.05; **, *p*<0.01. Scale bar: 50μm. **D**,**E,** P6 retinal vessels labelled with IB4 (grey) and stained for cleaved caspase 3 (red) in *YapTaz* iEC-KO (**E**) and littermate control mice (**D**).**D’**, **E’**, magnification of boxed area in D,E. Red arrowheads, cleaved caspase 3 positive endothelial cell. Black arrowheads, cleaved caspase 3 outside vessels. **D”**,**E”**, mask of cleaved caspase 3 positive endothelial cells. **F**, quantification of endothelial apoptosis in *Yap* iEC-KO (n=7 control/7 KO pups), *Taz* iEC-KO (n= 4 control/4KO pups) and *YapTaz* iEC-KO (n=5 control/4 KO pups). Data are mean +/- SD. *p* values were calculated using unpaired *t*-test. ns, *p*>0.05. Scale bar: D-E 100μm, D’-E’ 50μm. **G**, Quantification of endothelial proliferation with increasing concentrations of VEGF treatment in YAP, TAZ and YAP/TAZ knockdown cells and control. HUVECs were treated with 0, 4, 20 or 100 ng/mL VEGF for 24h and the percentage of cells in S phase was determined by flow cytometry. Graph shows the mean + SD fold change in percentage of S phase positive cells relative to 0 ng/mL of VEGF treatment. n= 3 independent experiments; > 50.000 cells analysed per experiment per condition. **H**, Quantification of endothelial proliferation after stretch in in YAP, TAZ, YAP/TAZ and VE-Cadherin knockdown cells and control. HUVECs were subjected to cyclic stretch for 24h and percentage of cells in S phase was determined by EdU pulsing and immunofluorescence staining. Graph shows the mean + SD fold change in percentage of S phase positive cells of stretched to non stretched cells for each knockdown condition. n= 5 independent experiments, > 100 cells counted per experiment per condition. *p* values were calculated using unpaired *t*-test. ns, *p*>0.05; *, *p*<0.05.

To understand if YAP and TAZ were required for proliferation downstream of VEGF, we knocked down YAP and TAZ in human umbilical vein endothelial cells (HUVECs) using small interfering RNAs (siRNAs) (Figure 3 - Figure supplement 1) and measured the proliferation rate by flow cytometry after treatment with increasing concentrations of VEGF (Figure 3G). Interestingly, upon loss of YAP, TAZ or YAP/TAZ, ECs proliferated at similar or even increased rates compared to control cells. Furthermore, VEGF treatment did not alter the subcelular localisation of YAP and TAZ in HUVECs (Figure 3 – Figure supplement 2), suggesting that VEGF is not a primary regulator of their activity.

We next asked whether YAP and TAZ mediate endothelial proliferation in response to stretch – another crucial mitogenic stimulus for the endothelium (19). To this end, we subjected HUVECs to 24h of stretch and measured the proliferation rate in comparison to non-stretched, static cells treated with the same siRNAs, by EdU labelling (Figure 3H). Control cells responded to stretch with a 5-fold average increase in proliferation, and this effect was reduced upon knockdown of VE-Cadherin confirming previous observations (19). The knockdown of YAP and YAP/TAZ, but not TAZ alone, led to a decrease in stretch induced proliferation. Thus YAP is, similarly to VE-cadherin, required for endothelial cell proliferation in response to mechanical stimulation at cell-cell junctions.

### YAP/TAZ loss leads to irregular endothelial cell distribution and haemorrhages

Further analysis of *Yap/Taz* iEC-KO retinas revealed severe defects at the sprouting front. *Yap/Taz* iEC-KO mutant retinas had 23% (+/- 12.3, *p*= 0.0113) fewer angiogenic sprouts than the control (Figure 4A,B yellow asterisks and Figure 4 - Figure supplement 1). Moreover, whereas control sprouts were elongated and showed long cellular protrusions towards the non vascularised front (Figure 4A’), sprouts in *Yap/Taz* iEC-KO retinas were rounder and lacked protrusions (Figure 4B’). The defective sprout morphology correlated with irregular spacing and frequent aggregations of ECs within the sprouts (Figure 4B’), arguing that migration and/or the rearrangement of ECs are perturbed in *Yap/Taz* mutant vessels. Additionally, the *Yap/Taz* iEC-KO vasculature displayed aberrant vessel crossings (Figure 4C,C’,C”,D,D’,D”), suggesting that vessels may frequently have failed to anastomose or stabilize connections following sprouting, and instead passed each other. Interestingly, defects in cellular rearrangements, sprouting elongation and anastomosis have previously been associated with altered stability or dynamics of endothelial cell junctions (20-24). The defects in morphology were coupled to defects in function as *Yap/Taz* iEC-KO retinas displayed large haemorrhages from vessel sprouts at the angiogenic front (Figure 4E,E’,F,F’), indicating loss of junctional integrity.

**Figure 4.**
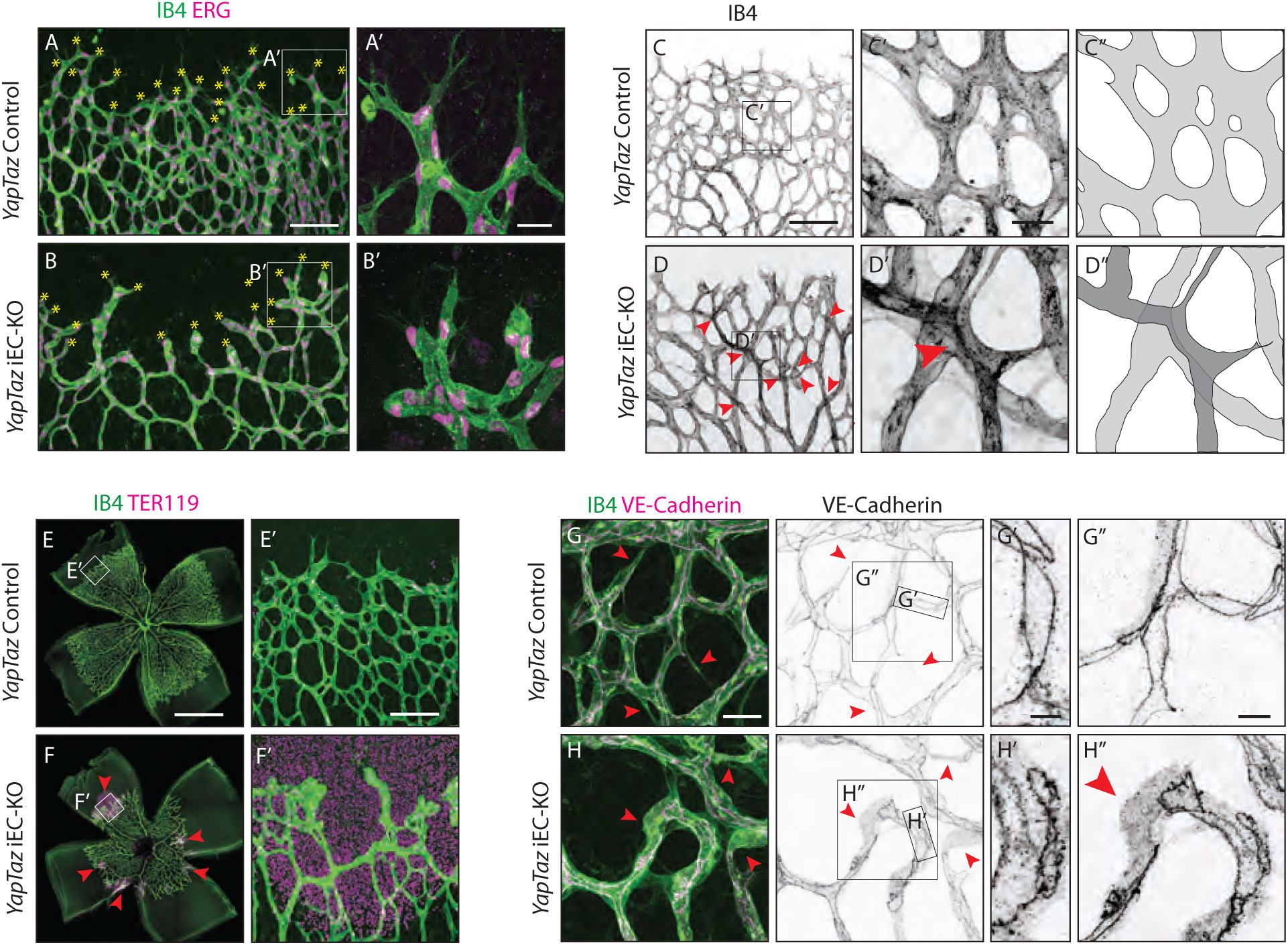
Combined loss of YAP and TAZ leads to decreased sprouting numbers and shape defects, vessel crosses, haemorrhages at the sprouting front and adherens junctions’ defects *in vivo.* **A,B**, P6 retinal vessels labelled with IB4 (green) and stained for ERG (magenta, marking endothelial nuclei) in *YapTaz* iEC-KO (B) and littermate control mice (A). Yellow asterisks mark sprouts. A’,B’, magnification of boxed areas in A and B. n=9 control/9 KO pups. Scale bar: A,B 100μm, A’, B’ 25μm. **C,D,** P6 retinal vessels labelled with IB4 in *YapTaz* iEC-KO (D) and littermate control mice (E). Red arrowheads, vessel crosses. C’, D’, magnification of boxed areas in C,D. C”,D”, depiction of vessels in C’ and D‘; different colours represent vessels in different 3D planes. n=4 control/4 KO pups. Scale bar: C,D 100μm, C’-D’ 20μm. **E,F**, P6 retinal vessels labelled with IB4 (green) and stained for TER119 (magenta, marking red blood cells) in *YapTaz* iEC-KO (F) and littermate control mice (E). Red arrowheads, haemorrhages. E’,F’, magnification of boxed areas in E and F. n=4 control/5 KO pups. Scale bar: E,F 1000μm, E’, F’ 100μm. **G,H**, P6 retinal vessels labelled with IB4 (green) and stained for VE-Cadherin (magenta) in *YapTaz* iEC-KO (H) and littermate control mice (G). Red arrowheads, no longitudinal VE-Cadherin labelled junction along vessel axis denoting unicellular vessel segments. G’,H’, G”,H”, magnification of boxed areas in G and H. n=4 control/4 KO pups. Scale bar: G,H 25μm, G’,H’ 5μm, G”,H” 10μm.

Together, these results argue against a cell proliferation defect being the only driver of the *Yap/Taz* iEC-KO phenotype and suggest that endothelial YAP/TAZ play a role in the regulation of ECs junctions.

### YAP/TAZ regulate adherens junction morphology and stability

Staining for VE-Cadherin revealed several junctional alterations in the vessels of *Yap/Taz* iEC-KO mice (Figure 4G-H). In control retinas, cell junctions were thin and mostly linear (Figure 4G’), while in *Yap/Taz* iEC-KO retinas ECs displayed tortuous junctions (Figure 4H’). VE-Cadherin staining also unveiled profound differences in the arrangement of ECs within vessels. In control retinas, ECs were arranged into multicellular tubes, highlighted by the presence of two or more VE-Cadherin junctions running longitudinally along the axis of the vessels (Figure 4G,G”). Some unicellular segments lacking VE-Cadherin staining could also be found and always correlated with decreasing calibre, indicative of regressing vessels (Figure 4G red arrowheads) (25). In contrast, in *Yap/Taz* iEC-KO retinas we observed many unicellular vessel segments lacking longitudinal VE-Cadherin junctions, but in vessels of normal calibre (Figure 4H, red arrowheads and H”). As junctional remodelling has been shown to be required for the cellular rearrangements that establish multicellular tubes (20), these results suggest that YAP and TAZ regulate junctional remodelling.

VE-Cadherin staining in HUVECs after YAP, TAZ and YAP/TAZ knockdown revealed altered junctional morphology. Interestingly, previous studies have correlated junctional morphology with cellular activities. *In vivo*, straight or linear junctions were associated with high Notch activity and stalk cell behaviour, while serrated junctions (also referred to as VE-Cadherin fingers) were found in tip cells or actively rearranging cells (21). *In vitro*, VE-Cadherin fingers were shown to steer migrating ECs and couple leader and follower cells (26), and have also been correlated with increased permeability in cell monolayers. More recently, junction associated intermediate lamellipodia have been associated with decreased permeability in cultured ECs (27).

To more accurately describe the differences in junctional morphology after YAP/TAZ knockdown, we defined five junctional categories (straight junctions, thick junctions, thick to reticular junctions, reticular junctions and fingers) (Figure 5E). Control cells showed mostly reticular junctions (Figure 5A,F). The knockdown of YAP and TAZ led to an increase in straight junctions and fingers, respectively (Figure 5B,C,F), whereas the combined knockdown of YAP/TAZ led to an increase in both straight junctions and fingers and to a loss of reticular junctions (Figure 5D,F). In addition, the knockdown of YAP/TAZ led to junctional breaks in the monolayer, as seen by the presence of gaps in VE-Cadherin stainings (Figure 5D, red arrowheads). Together, these observations demonstrate that YAP and TAZ together are required for the formation of reticular junctions and inhibit the formation of straight junctions and fingers. Interestingly, however, they individually have distinct effects on adherens junction morphology.

**Figure 5.**
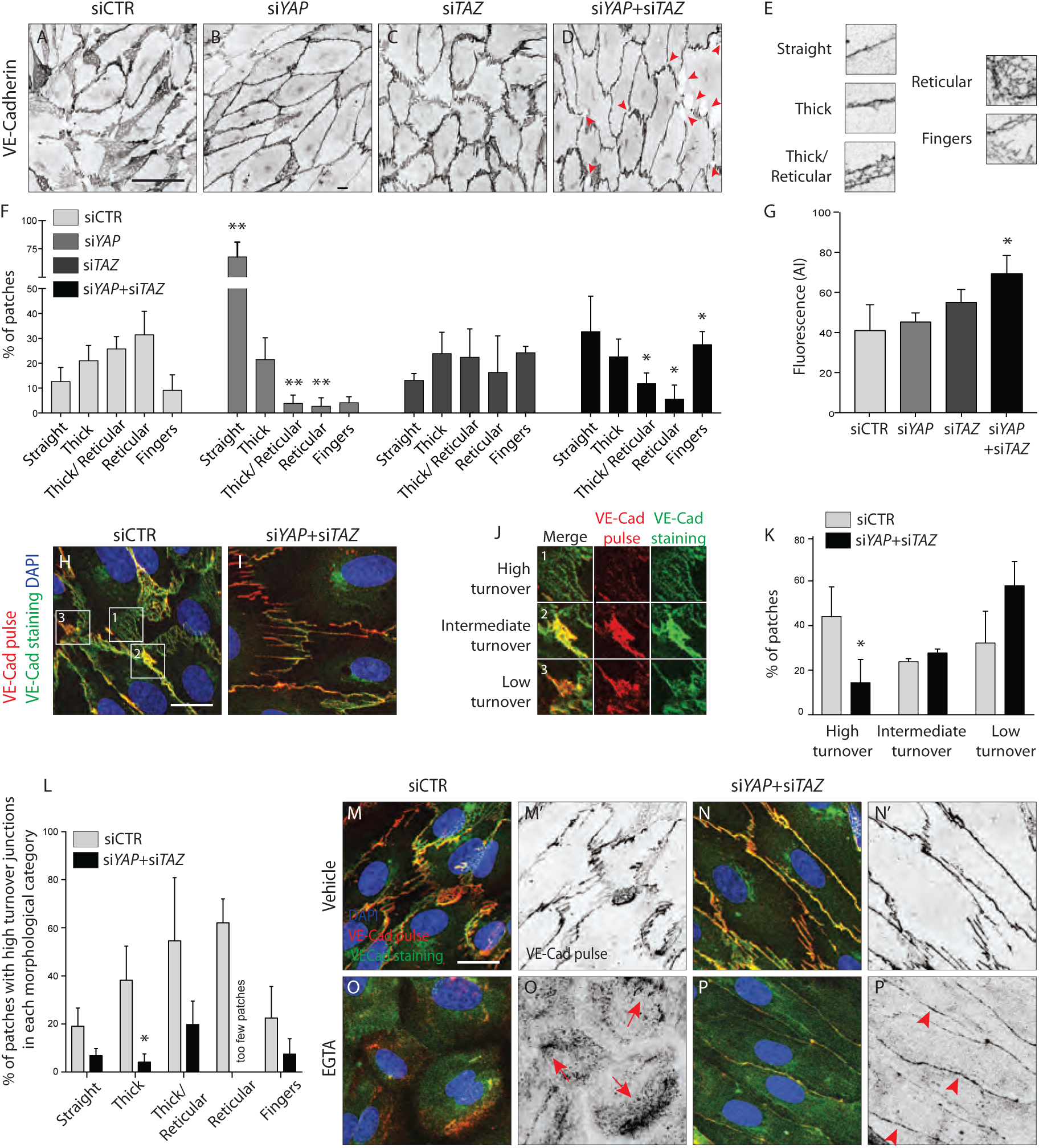
YAP and TAZ regulate adherens junctions’ morphology, monolayer permeability and VE-Cadherin turnover *in vitro*. **A-D**, HUVECs knocked down for YAP (B), TAZ (C) and YAP/TAZ (D) and control (A) stained for VE-Cadherin. Red arrowheads, discontinuous VE-Cadherin. Scale bar: 50μm. **E**, Representative patches used for manual morphological classification of adherens junctions in 5 categories: straight junctions, thick junctions, thick to reticular junctions, reticular junctions and fingers. **F**, Morphological analysis of VE-Cadherin labelled cell junctions in HUVECs knocked down for YAP, TAZ and YAP/TAZ. Data are mean percentage + SD of 3 independent experiments (2 for si*TAZ*). n> 140 patches of VE-Cadherin stained HUVECs per knockdown condition per experiment. *p* values were calculated using unpaired *t*-test between knocked down cells for YAP, TAZ and YAP/TAZ and control. *, *p*<0.05; **, *p*<0.01. **G**, Permeability of YAP, TAZ and YAP/TAZ knockdown monolayers of HUVECs to 250kDa fluorescent dextran molecules. Data are mean + SD of 3 independent experiments. *p* values were calculated using unpaired *t*-test between knocked down cells for YAP, TAZ and YAP/TAZ and control. *, *p*<0.05. **H**, **I**, HUVECs knocked down for YAP/TAZ (I) and control (H) triple labelled with DAPI (blue), pulsed VE-Cadherin 55-7HI (red, VE-Cadherin pulse), and surface VE-Cadherin (green, VE-Cadherin staining). VE-Cadherin 55-7HI pulse was done for 30 minutes and cells were fixed 2 hours after end of pulse. Scale bar: 20μm. **J**, Representative patches used for manual classification of junctions into high, intermediate and low turnover. **K**, Quantification of junctional turnover in YAP/TAZ knockdown cells and control. **L**, Quantification of the percentage of high turnover junctions in each morphological category in YAP/TAZ knockdown cells and control. **K**, **L**, Data are mean + SD of 3 independent experiments. n> 70 patches per knockdown condition per experiment. Fewer then 5 patches were reticular in YAP/TAZ knockdown, not allowing for reliable assessment of percentages between high, intermediate and low turnover. *p* values were calculated using unpaired *t*-test. *, *p*<0.05. **M-P**, HUVECs knocked down for YAP/TAZ (N,N’P,P’) and control (M,M’,O.O’) triple labelled with DAPI (blue), pulsed VE-Cadherin 55-7HI (red, VE-Cadherin pulse), and surface VE-Cadherin (green, VE-Cadherin staining) after treatment with EGTA (O,O’,P,P’) or vehicle (M, M’,N,N’). VE-Cadherin 55-7HI pulse was done for 30 minutes and cells were fixed 30 minutes after end of pulse, during which time were incubated with EGTA or vehicle. Red arrows, intracellular accumulation of pulsed VE-cadherin. Red arrowheads, pulsed VE-Cadherin at the junction. Scale bar: 20μm.

To understand whether the shift in morphology translated into a functional defect we investigated the permeability of the monolayer to 250kDa dextran molecules. Only the combined knockdown of YAP/TAZ led to a significant increase in permeability in comparison to the control situation (Figure 5G), suggesting that YAP/TAZ are both required for the barrier function of the endothelium and can compensate for each other in this particular role.

The dynamic rearrangements of ECs during sprouting require that cell-cell junctions are constantly assembled, rearranged and disassembled. To understand whether YAP and TAZ regulate the turnover of cell junctions, we pulse-labeled VE-Cadherin molecules at cell junctions using an antibody directly coupled to a fluorescent dye for 30 minutes (Figure 5H-I) (28). The antibody was subsequently washed out and cells cultured for two more hours in normal conditions, before being fixed and stained for surface VE-Cadherin using a second fluorescent label. Comparing the two sequential VE-cadherin labels allowed us to distinguish junctions with high, intermediate and low turnover rates (Figure 5J). In control cells, 44% of patches were of high turnover junctions, 24% of intermediate turnover junctions and 32% of low turnover junctions (Figure 5K). The knockdown of YAP/TAZ significantly decreased the percentage of high turnover junctions to 14% (*p*=0.0387) and increased the percentage of low turnover junctions to 58%. Interestingly, we found a correlation between the morphology of junctions and VE-Cadherin turnover rates (Figure 5L): straight junctions and fingers showed the lowest turnover rate, while reticular junctions showed the highest. To understand if the different VE-Cadherin turnover observed after knockdown of YAP/TAZ was caused by a shift in morphology, we compared the turnover of VE-Cadherin within the same morphological categories. Knockdown of YAP/TAZ decreased the percentage of high turnover junctions within all morphological categories, confirming a specific defect in VE-Cadherin turnover.

To test whether YAP and TAZ regulate the endocytosis of VE-Cadherin, we imaged intracellular VE-Cadherin vesicles after pulse labeling the molecule at the surface and allowing it to be endocytosed (Figure 5M,M’,N,N’). However, the detectable amount of VE-cadherin vesicles after YAP/TAZ knockdown was unaffected. In order to challenge the stability of the junctions we triggered cell junction disruption by chelating extracellular calcium with EGTA (29) (Figure 5O,O’,P,P’). Whereas VE-cadherin accumulated in intracellular vesicles in control cells, after YAP/TAZ knockdown substantial amounts of VE-cadherin antibody remained at the cell junction, signifying reduced endocytosis. Thus, YAP/TAZ promote VE-Cadherin turnover and facilitate its dynamic recycling.

As cell junctions are essential for ECs to rearrange and migrate collectively (30), and *Yap/Taz* iEC-KO retinas presented less elongated sprouts suggestive of a migration defect, we asked whether cell migration was also regulated by YAP/TAZ.

### YAP/ TAZ are required for individual endothelial cell migration

To address the requirement of YAP and TAZ for endothelial cell migration we performed a scratch-wound assay (Figure 6A-H). While in the control situation the wound was completely closed at 16h (Figure 6A,B,I), after knockdown of YAP less than 50% of the wound area was closed at the same time point (*p*=0.0067) (Figure 6C,D,I). A stronger effect on endothelial cell migration was observed after the knockdown of TAZ and YAP/TAZ, with less than 20% of the wound area being closed at 16h (*p*=0.0006 for siTAZ vs siCTR and *p*=0.0013 for siYAP+siTAZ vs siCTR) (Figure 6E,F,G,H,I).

**Figure 6.**
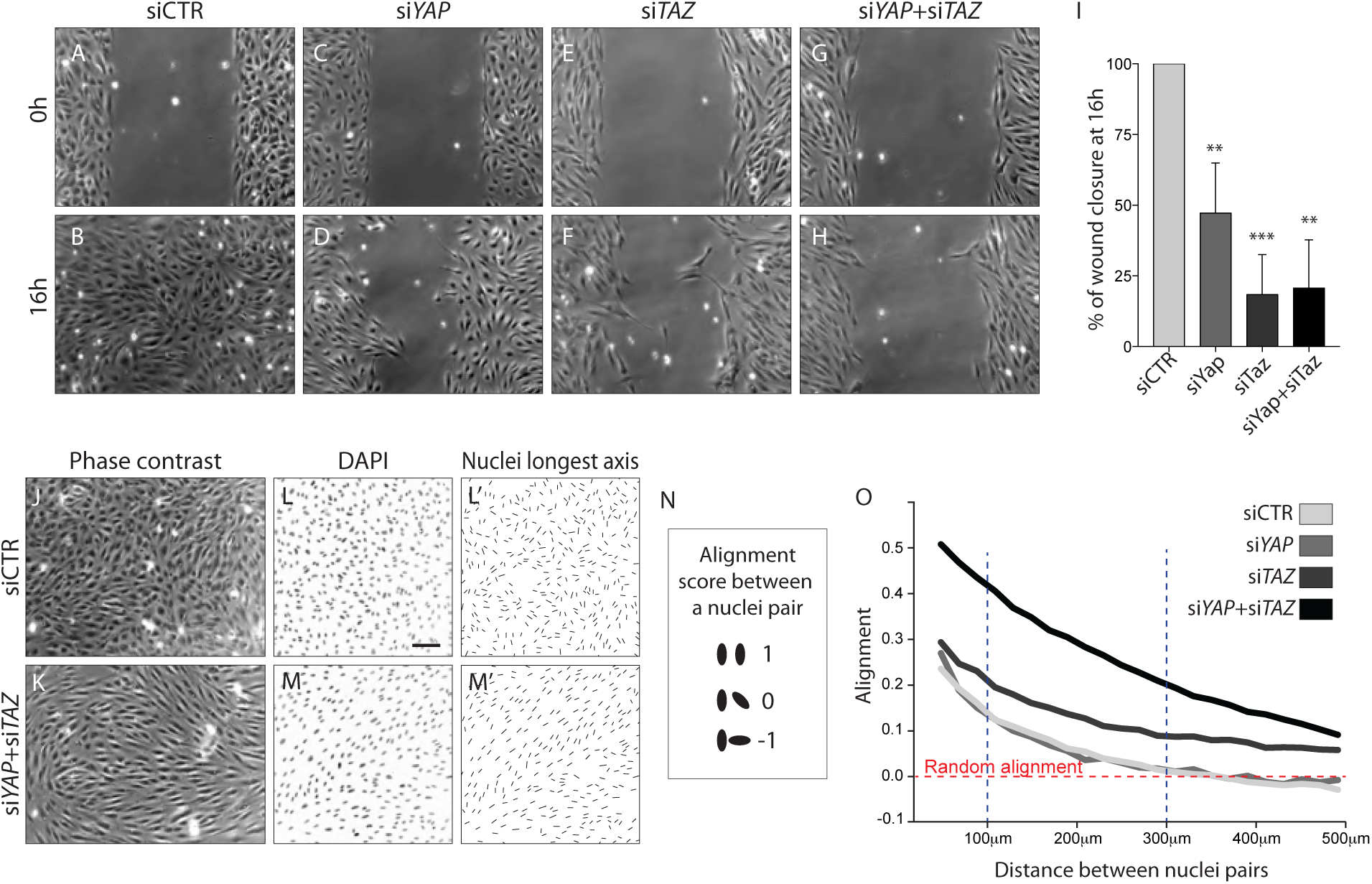
YAP and TAZ are required for uncoupled, individual cell migration. **A-H**, Phase contrast images of YAP (C,D), TAZ (E,F) and YAP/TAZ (G,H) knockdown HUVECs and control (A,B) immediately after removing barrier to create a cell free space (A,C,E,G) and 16 hours later (B,D,F,H). **I**, Quantification of wound closure at 16 hours. Data are mean + SD of 3 independent experiments. *p* values were calculated using unpaired *t*-test between knocked down cells for YAP, TAZ or YAP/TAZ and control. **, *p*<0.01; ***, *p*<0.001. **J,K**, Phase contrast images of YAP/TAZ knockdown monolayer of HUVECs (K) and control (J). **L,M**, Fluorescence labelling of nuclei with DAPI of YAP/TAZ knockdown monolayer of HUVECs (M) and control (L). Scale bar: 100μm. **L’,M’**, Longest axis of nuclei. **N**, Alignment score between nuclei pairs used for quantification of cell coordination in O. Angles made by the nuclei longest axis of a pair of nuclei were calculated; angles of 0, 45 and 90 degrees scored 1,0 and -1 in alignment. **O**, Coordination plot of monolayers of HUVECs knocked down for Yap, Taz and Yap and Taz and control. Graph shows mean alignment score of all pairs of cells in the monolayer plotted against distance between them. Randomly aligned cells score 0 in mean alignment. n= 3 independent experiments, >10.000 pairs of nuclei analysed per knockdown condition per experiment.

Given that cells aggregated at the sprouting front of *Yap/Taz* iEC-KO retinas, we wondered whether in addition to defective directional cell migration they also lacked the ability to shuffle with the neighbouring cells. Recent data illustrated that collectively migrating ECs *in vitro* move in streams and swirls and display straight junctions along the lateral boundaries and fingers along the front and rear (26). To investigate collective cell migration we therefore analysed the arrangement of cells in a confluent monolayer (Figure 6 J-O). Control cells displayed a cobblestone appearance without identifiable subgroups of cells (Figure 6J). In contrast, after knockdown of YAP/TAZ cells adopted elongated shapes and arranged into streams and swirls (Figure 6K). To quantity this effect we used the longest axis of the EC nucleus as a proxy for the orientation of each cell and developed a measure of monolayer coordination based on the alignment of cells with their neighbours (Figure 6L,L’,M,M’, N). A score of 1 would signify parallel alignment between all cells, and a score of 0 random alignment of the population. Control cells displayed higher than random alignment with their closest neighbours, but cells beyond 300μm from each other were arranged at random (Figure 6I). While the knockdown of YAP did not affect the alignment score of cells, the knockdown of TAZ led to increased alignment. The combined knockdown of YAP/TAZ led to an even higher degree of coordination, with higher alignment scores across all distances between cells. These results suggest that YAP/TAZ promote the ability of cells to distribute individually within monolayers.

### Nuclear YAP and TAZ inhibit Notch and BMP signalling in endothelial cells

To gain insight into the nuclear function of YAP and TAZ, we generated a *Pdgfb-iCreERT2* -inducible TAZ gain-of-function mouse allele, in which a mutated version of TAZ (TAZ S89A) is introduced in the Rosa26R locus and expressed by a CAG promoter following Cre-mediated excision of an upstream stop codon (Figure 7 – Figure supplement 1). The TAZ S89A mutation results in enhanced nuclear localization of TAZ as it escapes phosphorylation by the upstream Hippo kinase LATS (31). The allele also expresses nuclear EGFP by means of an IRES sequence, allowing the identification of recombined cells expressing the TAZ mutant protein. *Taz* iEC-GOF retinas exhibited 25% increased sprouting (+/- 12.2, *p*=0.0074) (Figure 7A,B yellow asterisks and Figure 7C) and 19% increased branching (+/- 8.3, *p*=0.0012) (Figure 7D). Thus, driving nuclear TAZ expression, leads in many aspects to the opposite phenotype of *Yap/Taz* iEC-KO retinas, suggesting that the loss of the nuclear function of YAP/TAZ plays a key role in the development of the observed vascular loss-of-function phenotypes.

**Figure 7.**
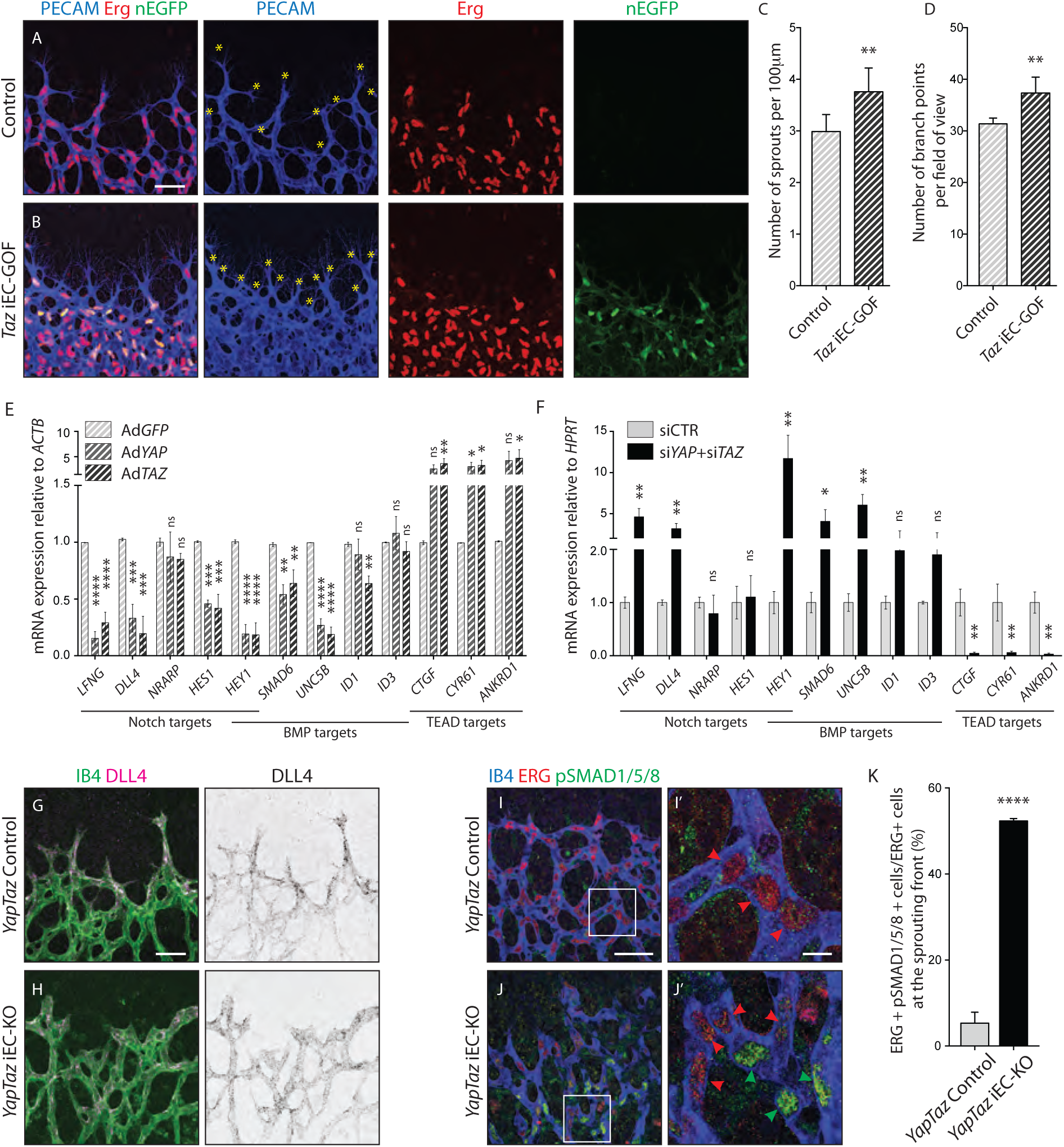
Nuclear YAP and TAZ inhibit Notch and BMP signalling in endothelial cells. **A-B**, Retinas from P6 *Taz* iEC-GOF (B) and control pups (A) were stained for the endothelial marker PECAM (blue) and the endothelial nuclei marker ERG (red). *Taz* iEC-GOF mice express mosaically nuclear EGFP (nEGFP, green) marking cells expressing the TAZ gain of function mutation TAZS89A. Yellow asterisks mark sprouts. Images correspond to maximum projection of z stack. Scale bar: 50μm. **C**, Quantification of number of sprouts per 100 μm of sprouting front extension at P6 in *Taz* iEC-GOF mice (n= 6 pups) and littermate control mice (n=6 pups). Data are mean + SD. *p* values were calculated using unpaired *t*-test. **, *p*<0.01. **D**, Quantification of branching frequency (i.e. number of branching points per field of view) in *Taz* iEC-GOF mice (n= 6 pups) and littermate control mice (n=6 pups). Data are mean + SD. *p* values were calculated using unpaired *t*-test. **, *p*<0.01. **E**, Reverse transcriptase PCR of HUVECs transduced with adenoviruses carrying *YAP* (Ad*YAP*) and *TAZ* (Ad*TAZ*) constitutively active forms and control (Ad*GFP*). Data are mean + SD of 3 independent experiments. *p* values were calculated using one-way ANOVA. *, *p*<0.05; **, *p*<0.01; ***, *p*<0.001; ****, *p*<0.0001. **F**, Reverse transcriptase PCR of YAP/TAZ knockdown HUVECs and control. Data are mean + SD of 3 independent experiments. *p* values were calculated using unpaired *t*-test. *, *p*<0.05; **, *p*<0.01; ***, *p*<0.001; ****, *p*<0.0001. **G,H**, P6 retinal vessels labelled with IB4 (green) and stained for DLL4 (magenta) in *YapTaz* iEC-KO mice (H) and littermate control mice (G). Images correspond to maximum projection of z stack. Scale bar: 50μm. **I,J**, P6 retinal vessels labelled with IB4 (blue) and stained for ERG (red, marking endothelial nuclei) and pSMAD1/5/8 (green) in *YapTaz* iEC-KO (J) and littermate control mice (I). Images correspond to single confocal planes. I’,J’, magnification of boxed areas in I and J. Red arrowheads, endothelial nuclei negative for pSMAD1/5/8. Green arrowheads, endothelial nuclei positive for pSMAD1/5/8. Scale bar: I,J 50μm, I’, J’ 10μm. **K**, Quantification of endothelial cells positive for pSMAD1/5/8 at the sprouting front of the P6 retina in *YapTaz* iEC-KO (n= 3 pups) and littermate control mice (n=3 pups). Data are mean percentage + SD. *p* values were calculated using unpaired t-test. ****, p<0.0001.

To elucidate the transcriptional targets of YAP and TAZ, we performed unbiased transcriptome analysis on HUVECs transduced with adenoviruses encoding for *YAP* and *TAZ* gain-of-function mutants or *GFP* as a control (AdYAP^S127A^, AdTAZ^S89A^, AdGFP) (Figure 7 – Figure supplement 2). Forced activation of YAP and TAZ led to congruent gene expression changes including the canonical YAP/TAZ target genes *CYR61*, *ANKRD1*, and *CTGF*, as expected. Interestingly, YAP and TAZ also suppressed numerous Notch and BMP target genes. During sprouting angiogenesis, Notch and BMP9/10 signalling restrict the acquisition of a tip cell phenotype by activated ECs (32-38). These results were confirmed by qRT-PCR analysis (Figure 7E): AdYAP^S127A^ and AdTAZ^S89A^ cells expressed significantly less *LFNG*, *DLL4* and *HES1* (Notch target genes), *SMAD6*, *UNC5B* and *ID1* (BMP target genes) and *HEY1* (a common Notch and BMP target gene) than control cells. Consistent with these findings, knockdown of YAP or TAZ lead to a substantial increase in Notch reporter activity (Figure 7 – Figure supplement 3A) and target gene expression (Figure 7F). Similar effects were observed for the BMP pathway (Figure 7 – Figure supplement 3B and Figure 7F), while TEAD-driven reporter activity and YAP/TAZ target genes were repressed (Figure 7 – Figure supplement 3C and Figure 7F).

To understand if Notch and BMP signalling were also affected *in vivo*, we stained *Yap/Taz* iEC-KO retinas for DLL4 and phospho-SMAD1/5/8. In control retinas, DLL4 expression was highest at the leading edge, decreasing over the first 100μm from the sprouting front, beyond which the expression was evenly low throughout the vessels in the plexus (Figure 7G,H and Figure 7 – Figure supplement 4). In *Yap/Taz* iEC-KO retinas the expression of DLL4 at the sprouting front was higher; additionally, the area of high DLL4 was broader, decreasing for up to 200μm from the sprouting front before flattening to the lower levels of the plexus. Moreover, staining *Yap/Taz* iEC-KO retinas for pSMAD1/5/8 showed a ∼10 fold increase in the number of ECs positive for pSMAD1/5/8 at the sprouting front (*p*<0.0001) (Figure 7,I’, J, J’,K).

Together, these results identify that endothelial YAP and TAZ repress Notch and BMP signalling during angiogenesis and retinal vascular expansion.

## DISCUSSION

The present study aimed to provide a detailed understanding of the distribution and function of endothelial YAP and TAZ in angiogenesis. Our finding that YAP and TAZ were present in the nucleus of ECs at the sprouting front of developing vessels shows parallels with other cell types where nuclear YAP and TAZ are detected in actively proliferating areas of developing tissues. Interestingly, however, YAP and TAZ show distinct expression patterns in ECs, although these proteins show a high degree of redundancy in many other cell types. While TAZ was predominantly expressed in the sprouting front where it accumulated strongly in endothelial nuclei, YAP was mostly cytoplasmic both in the sprouting front and also in more mature, remodelling vessels. These data are in agreement with a role for YAP in vessel maintenance. Also in zebrafish, the loss of YAP did not impact vessel formation but led to vessel collapse and regression (14). Furthermore, in addition to nuclear and cytoplasmic YAP and TAZ, we also detected junctional localization of these proteins in retinal vessels. A previous study by Giampietro and colleagues (7) has shown that endothelial YAP associates with adherens junction proteins at stable junctions and that this prevents its nuclear accumulation and transcriptional activity. Whether this is also true for TAZ has previously not been addressed. A sequestration of YAP and TAZ either in the cytoplasm or bound to junctional proteins can potentially serve different and not necessarily mutually exclusive roles: preventing their nuclear activity, keeping a pool of protein ready to shuttle to the nucleus and drive gene expression, and having other cytoplasmic functions. It is not yet entirely clear what regulates the subcellular localisation of YAP and TAZ in the developing vasculature. ECs at the sprouting front and in more mature vessels have different adherens junctions, experience distinct levels of signalling from secreted angiogenic molecules and are exposed to different levels of shear stress by the blood. Adherens junctions in sprouting vessels undergo dynamic remodelling that accompanies endothelial cell shape changes and migration, in contrast to mature vessels where they appear more stable in terms of shape (21, 39). Endothelial YAP and TAZ relocate to the nucleus upon disruption of cell junctions or loss of VE-Cadherin (shown for YAP by Choi and colleagues (16) and confirmed in our analysis also for TAZ, data not shown). Interestingly, we did not find junctional localisation of YAP or TAZ at the sprouting front, supporting the idea that more dynamic junctions fail to sequester YAP and TAZ away from the nucleus. Conceptually, these data would support the hypothesis that the subcellular localisation of YAP and TAZ in the vasculature is at least in part regulated by the maturation of adherens junctions. In zebrafish, Nakajima and colleagues (14) showed that YAP nuclear relocation correlated with lumenisation of sprouting vessels, and they attributed this to the effect of shear stress on YAP. In the mouse retina, hemodynamic fluid laws predict that vessels at the sprouting front experience very low levels of shear (40), arguing against YAP and TAZ being activated by shear in this model. Additionally, we found no difference in the subcellular localisation of YAP or TAZ between arteries and veins, i.e. vessels that experience distinct shear stress levels. However, it is possible that local and fast changes in shear stress levels are more relevant to regulate YAP and TAZ than sustained shear. In support of this idea, YAP and TAZ appear not to respond to 12 or 24h of laminar shear (41), but translocate to the nucleus after only 10 minutes of laminar shear (14). Finally, although VEGF, a pro-angiogenic molecule secreted by astrocytes at the avascular front, drives endothelial proliferation and migration, we found no evidence for VEGF induced YAP and TAZ nuclear translocation. Other pro-angiogenic molecules, either locally produced or blood-borne, could regulate endothelial YAP and TAZ during development; future work will help clarify these questions and how different chemical and mechanical stimuli come together to regulate YAP and TAZ.

The role of YAP in the development of the retinal vasculature has previously been studied by Choi and colleagues using intra-ocular injection of siRNAs (16). However, we observed that YAP and TAZ are also expressed in pericytes. Thus to address the cell autonomous role of YAP and TAZ we took advantage of an endothelial specific inducible Cre to inactivate YAP and/or TAZ genetically during angiogenesis. The mild phenotype of the single mutants in comparison to the drastic phenotype of the compound mutant indicates functional redundancy in the endothelium. The compound loss of endothelial YAP and TAZ leads in the mouse retina to a decrease in the radial expansion of vessels, vascular density, branching and sprouting. This phenotype could be a consequence of a decreased number of ECs caused by a proliferation defect (42). However, our further discovery that YAP and TAZ are required to establish homogeneity in the plexus and prevent cellular aggregations suggests that endothelial YAP/TAZ signalling is not only required to provide adequate number of cells but is also critically involved in ensuring adequate EC distribution. We propose that endothelial YAP/TAZ operate in several mechanisms that jointly establish a balance of the right number of endothelial cells in the right place. First, endothelial YAP/TAZ drive proliferation in response to mechanical stimulation at the cell-cell junction, and not in response to VEGF. We propose that in this way endothelial YAP/TAZ provide a cell intrinsic mechanism of locally controlling cell densities, in contrast to growth factor mediated cell proliferation instructed by the surrounding tissue. Second, endothelial YAP/TAZ increase VE-Cadherin turnover at cell-cell junctions, which is essential for cells to rearrange. This corroborates recent findings in mouse hepatocytes where YAP antagonises adherens junction stability (43). The authors showed that YAP regulates hepatocyte adherens junctions in response to increased actomyosin contractility by increasing myosin II light chain gene expression. Accordingly, the transcriptional, nuclear role of YAP was required for junctional regulation. Together with our observations, these findings indicate the existence of a positive feedback loop where stable junctions sequester YAP and TAZ from the nucleus, therefore maintaining less junctional turnover, while remodelling junctions allow YAP and TAZ to relocate to the nucleus where they increase VE-Cadherin turnover. Our results also suggest that a high VE-Cadherin turnover at the sprouting front is required in order to maintain junctional integrity and prevent bleedings. Third, YAP/TAZ decrease cell-cell coupling and increase the ability of cells to migrate individually. Together, our results therefore identify a role for YAP/TAZ in promoting endothelial cellular rearrangements through the regulation of junctional turnover and collectiveness of cell migration. Conceptually, linking stretch induced proliferation to balance cell numbers with modulation of junctional turnover to facilitate cell rearrangements seems ideally suited to achieve the required balance of cell distribution for functional vascular patterning.

Molecularly, how YAP and TAZ affect this complex cell behavior is not clear. Our results identify that endothelial YAP/TAZ reduce the expression of Notch and BMP signaling in ECs. Interestingly, temporal fluctuations of Notch signalling in sprouting ECs are required for their shuffling behaviour in sprouting assays, and heterogeneity in the Notch-activation phase between contacting cells drives vessel branching (42). Given that YAP/TAZ dynamically shuttle between cytoplasm and nucleus, it is tempting to speculate that they may affect not only Notch-signalling levels, but also their dynamics. However, further work and new tools will be required to address these questions. On the other hand, the BMP9/10-Alk1 pathway has recently been proposed as an important driver of collective migration of ECs in particular in response to blood flow (44). Therefore, the newly identified roles of endothelial YAP/TAZ in regulating Notch and BMP signalling may prove critical for endothelial cell migration and rearrangements not only within new vascular sprouts, but also in already perfused vessels. Based on our current evidence, we propose that endothelial YAP/TAZ function as integrators of mechanical stimulation and Notch/BMP signaling to balance local cell densities and endothelial cell arrangements during sprouting angiogenesis.

## MATERIAL AND METHODS

### Mice and treatments

For loss of function experiments the following mouse strains were used: *Yap* ^*fl/fl*^ and *Taz* ^*fl/fl*^(17), *Pdgfb-iCreERT2* (18). To generate a conditional TAZ gain-of-function mouse model, a cDNA coding for a 3xFLAG-tagged human *TAZ* (*WWTR1*) carrying an alanine substitution at serine 89 (S89A)(45) was inserted into a *Rosa26* targeting vector downstream of the ubiquitous *CAG* promoter. The cDNA also included an internal ribosome entry sequence (*IRES*) and a *nuclear-localized enhanced green fluorescence protein* (*nEGFP*)(46) for monitoring transgene expression. To allow Cre-dependent expression of *3xFLAG-TAZ*^*S89A*^ and of the EGFP reporter, a *floxed* (*loxP*-flanked) transcriptional *STOP* cassette was incorporated between the *3xFLAG-TAZ*^*S89A*^*-IRES-nEGFP* sequence and the *CAG* promoter. The linearized targeting vector was transfected into embryonic stem (ES) cells derived from C57BL/6 mice, and homologous recombinants were identified by Southern blotting analysis. Correctly targeted ES cells were implanted into foster mothers and resulting chimaeras bred to C57BL/6 mice to screen for germline transmission. The mouse model was developed together with genOway. The *Rosa26-3xFLAG-TAZ*^*S89A*^*-IRES-nEGFP* allele was kept heterozygous in the experimental studies.

Mice were maintained at the London Research Institute and at the Max Delbruck Center for Molecular Medicine (loss of function mice) and at the Max Planck Institute for Heart and Lung Research (gain of function mice) under standard husbandry conditions. To induce Cre-mediated recombination 4-hydroxytamoxifen (Sigma, 7904) was injected intraperitoneally (IP) (20 μL/g of 1 mg/mL solution) at postnatal day 1 and day 3 and eyes were collected at P6. In all loss and gain of function experiments control animals were littermate animals without Cre expression. Male and female mice were used for the analysis.

For endothelial cell proliferation assessment in the retina, mouse pups were injected IP 2 hours before culling with 20 uL/g of EdU solution (0.5 mg/mL; Thermo Fischer Scientific, C10340).

### Cell culture

HUVECs from pooled donors (PromoCell) were cultured in EGM2-Bulletkit without antibiotics (Lonza) and used until passage 6. For YAP and TAZ gain of function experiments HUVECs were obtained from Lonza, cultured in endothelial basal medium (Lonza) supplemented with hydrocortisone (1 μg ml^−1^), bovine brain extract (12 μg ml^−1^), gentamicin (50 μg ml-1), amphotericin B (50 ng ml^−1^), epidermal growth factor (10 ng ml^−1^) and 10% fetal bovine serum (Life Technologies) and used until passage 4.

For knockdown experiments, HUVECs were transfected with SMARTpool: siGENOME siRNAs purchased from Dharmacon (Yap #M-012200-00-0005, Taz #M-016083-00-0005, VE-Cadherin # M-003641-01-0005 and non-targeting siRNA Pool 1 #D001206-13-05). Briefly, subconfluent (70-80%) HUVECs were transfected with 25 nM siRNA using Dharmafect 1 transfection reagent following the protocol from the manufacturer; transfection media was removed after 24h and experiments were routinely performed on the third day after transfection.

To activate YAP and TAZ signalling in ECs, FLAG-YAP^S127A^- or 3x-FLAG-TAZ^S89A^-encoding adenoviruses were generated in the adenoviral type 5 backbone lacking the E1/E3 genes (Vector Biolabs). GFP-encoding adenoviruses were used as a control. Infections were carried out by incubating sub-confluent HUVECs (70-80%) with starvation media (EBM containing 0.1% BSA) for 4 hours followed by the addition of adenoviral particles and polybrene (Santa Cruz). After 4 hours, HUVECs were washed with Hanks Buffer for at least five times and then cultured in complete EBM media with 10% FCS and supplements overnight. All experiments were performed 24 hours post transduction.

### Immunofluorescence staining

To perform retina immunofluorescence, eyes were collected from postnatal day 6 mice and fixed in 4% PFA in PBS for 1h at 4C. Retinas were dissected in PBS and permeabilised/ blocked for 1h at room temperature in 1% BSA, 2% FBS, 0.5% Triton X100, 0.01% Na deoxycholate and 0,02% Na Azide in PBS. Primary and secondary antibodies were incubated overnight at 4C and for 2h at room temperature, respectively, both in 1:1 PBS: blocking buffer. Isolectin staining was performed overnight at 4C in Pblec after retinas were equilibrated for 1h in Pblec at room temperature. Retinas were post-stained fixed in 2% PFA in PBS for 10 minutes. To mount the samples Vectashield mounting medium. (Vector Labs, H1000) or ProLong Gold (Thermo Fisher Scientific) was used. Imaging was done by laser scanning confocal microscopy (Carl Zeiss LSM700, LSM780 and Leica TCS SP8). Processing of samples was carried out in tissues from littermates under the same conditions.

For immunofluorescence in HUVECs, cells were grown in #1.5 coverslips coated with polylysine and gelatin 0.2%. At the end of the experiment cells were fixed in 4% PFA for 10min, permeabilised in 0.3% Triton-X100 in blocking buffer for 5min and blocked in 1% BSA 20mM Glycine in PBS for 30 min. Primary and secondary antibodies were incubated for 2 and 1 hours, respectively, in blocking buffer. Nuclei labeling was performed by incubating cells with DAPI for 5 min (Life technologies, D1306).

A list of the primary antibodies used can be found in Supplementary Table 1.

### Image analysis

Analysis of radial expansion, capillary density, branching frequency, proliferating ECs, apoptosis and sprouting numbers was done using Fiji. Radial expansion corresponds to the mean distance from the optic nerve to the sprouting front (8 measurements in tilescans of two whole retinas per animal). Capillary density corresponds to the vessel area (measured by thresholding IB4 signal) divided by the field of view area (6–8 images of (425 μm)^2^ between artery and vein per animal). Branching frequency was measured by manually counting all branching points in a field of view (4-5 images of (200 μm)^2^ between artery and vein per animal). The plexus regularity was assessed through the standard deviation of the size and the circularity of the vascular loops in the plexus (using same images as for analysis of capillary density). Vascular loops were segmented by thresholding the IB4 signal to avoid artifacts we excluded loops with a size smaller than 86 um^2 for the analysis. Endothelial proliferation was measured by manually counting the number of EdU positive endothelial nuclei (ERG positive) and dividing by the vessel area (measured by thresholding IB4 signal) (4 images of (425 μm)^2^ containing the sprouting front and localized on top of arteries per animal). Apoptosis was measured manually by counting the number of cleaved caspase 3 positive figures and dividing by the vessel area (measured by thresholding IB4 signal) (tilescan of one whole retina per animal). The number of sprouts was measured manually (3 images of 425x850μm of the sprouting front per animal). To quantify DLL4 intensity the outline of the sprouting front and the position of the arteries were manually defined using IB4 staining. Vessels were segmented by thresholding the IB4 staining in Fiji. Then, DLL4 intensity inside the vasculature was normalised with the average DLL4 intensity outside of the vasculature. Subsequently, for every pixel inside the vasculature (excluding the arteries) the distance to the sprouting front was calculated. The normalised DLL4 values within each bin were averaged (15μm bins from 0 to 500 μm). For each retina quarter a curve was obtained, and the average and SEM of these curves was shown in the graph (one retina quarter was used per animal). To quantify pSMAD1/5/8 status the number of pSMAD1/5/8 positive endothelial nuclei was manually counted and dividing by the total number of endothelial nuclei (defined by being ERG positive) (3 images of (225μm)^2^ containing the sprouting front were used per animal).

To analyse YAP/TAZ subcellular localisation in HUVECs we adapted a previously existing cytoplasm-to-nucleus translocation assay pipeline from Cell Profiler (47). Briefly, YAP or TAZ staining intensity was measured both inside the nucleus of the cell and in a 12 pixels wide ring of cytoplasm grown radially from the nucleus. The nucleus localisation was determined using a DAPI mask. We calculated the ratio between the nucleus and cytoplasm intensity and categorised cells as having nuclear localisation, nuclear and cytoplasmic and cytoplasmic localisation when the ratio was >1.2, between 1.2 – 0.8 and <0.8 respectively.

Cell junction morphology analysis was done in confluent monolayers of HUVECs stained for VE-Cadherin. 5 morphological categories were defined: straight, thick, thick to reticular, reticular and fingers. We acquired 5 images of (160μm)^2^ per condition per experiment, divided each image in (16 μm)^2^ patches, and randomly grouped these patches. The classification into categories was done manually and blindly for the condition.

To analyse cell coordination we used confluent cells labelled for DAPI. The nuclei were automatically segmented using a customized Python algorithm relying on the Scikit Image Library. By fitting an ellipse to each nucleus we obtained its major and minor axis, and the angle of the major axis with the x-axis of the image was assigned to the nucleus as its orientation. This way each nucleus in the images was assigned a position given by its midpoint and an orientation. Next we analyzed the average alignment of the nuclei of two cells depending on their distance. As the nuclei don’t have a directionality (i.e. they are nematics as opposed to vectors), the angles between two nuclei range from 0 corresponding to the nuclei being parallel, to *π/2* corresponding to them spanning a right angle. For any two cells in each image we calculated the angle and the Euclidean distance between them, and then we binned the cells depending on their distance. We introduced a parameter called ‘alignment’ which is 1 if all cells are perfectly aligned and 0 for a completely random distribution of cell orientations.

### VEGF treatment and YAP/TAZ staining

Confluent HUVECs were maintained in VEGF free media for 24h. VEGF treatment was then performed for 6h with 0ng/mL, 4ng/mL, 20ng/mL or 100ng/mL of VEGF-165 (PrepoTech, 450-32). Immunofluorescence staining and analysis of YAP and TAZ subcellular localisation was performed as above described.

### VEGF treatment and proliferation assessment

Knockdown HUVECs were maintained in VEGF free media for 24h. VEGF treatment was then performed for 24h with 0ng/mL, 4ng/mL, 20ng/mL or 100ng/mL of VEGF-165. Cells were pelleted, ressuspended in 90% cold Methanol and stored at -20C° before further processing. Cells were then ressuspended in Propidium Iodide/RNase staining solution (Cell signaling, 4087) for 30 minutes before cell cycle analysis by flow cytometry (LSRII, BD). Data was analysed using BD FACSDiva^™^ software.

### Mechanical stretch application and proliferation assessment

HUVECs were plated on collagen I - 0.2% gelatine-coated Bioflex plates (BF-3001C, Flexcell International Corporation). Gene knockdown was preformed as previously described. Cells were incubated in transfection media for 24h, and allowed to recover in fresh complete media for 4h. Afterwards cells were incubated for 24h in serum starvation media (0,1%BSA in EBM2 pure media) to form a confluent, quiescent monolayer. Cyclic stretch (0.25Hz, 15% elongation) was then applied for 24h using a Flexcell® FX-5000^™^ Tension System. Control cells were placed in the same incubator but not on the Flexcell® device (static conditions). EdU pulsing was performed after 20h of the 24h stretch period. At the end of the experiment cells were fixed in 4% PFA and EdU staining was performed according to the manufacturer’s protocol (Click-It EdU C10337 Life Technologies). Nuclei were labelled with DAPI. Three regions of interested were acquired per sample in a Carl Zeiss LSM700 scanning confocal microscopes (Zeiss, Germany). Quantification of proliferation was done using a CellProfiler pipeline. Percentage of S phase cells was determined as percentage of EdU positive nuclei over the total number of nuclei.

### Permeability assay

24h after siRNA transfection cells were re-plated into fibronectin coated Transwell® membranes (Costar 3460) at confluence and incubated for 2 more days to stabilize cell junctions. On the third day after transfection 0.5mg/mL of 250kDa FITC Dextran in cell media (Sigma FD250) was added to the top well. Fluorescence on the bottom well was measured after 6h in a Gemini XPS fluorescent plate reader. In each experiment 3 wells were measured per condition.

### Pulse chase VE-Cadherin experiment for quantification of low, intermediate and high turnover junctions

Cells were labelled live with a non-blocking monoclonal antibody directed against extracellular VE-Cadherin and directly coupled with Alexa-Fluor647 (BD Pharmingen, #555661, 1:200) for 30 minutes. Cells were then washed 2x with PBS and incubated with complete media for additional 2 hours. Cells were fixed with 4% PFA and stained for VE-Cadherin (Santa Cruz Biotechnology, #6458, 1:200) with a secondary antibody coupled with Alexa-Fluor-488. 5 (160μm)^2^ images per condition per experiment were acquired in a Carl Zeiss LSM700 confocal laser scanning microscope using the same acquisition settings. Max projection of z stack and merging of channels was done in Fiji. Images were divided in (16 μm)^2^ patches and the patches were randomly grouped. Patches were classified into a morphological category and into low, intermediate or high turnover categories, manually and blindly for the condition.

### Pulse chase VE-Cadherin experiment and EGTA treatment

Cells were labelled live with a non-blocking monoclonal antibody directed against extracellular VE-Cadherin and directly coupled with Alexa-Fluor647 (BD Pharmingen, #555661, 1:200) for 30 minutes. Cells were then washed 2x with PBS and incubated with 4mM EGTA or vehicle in complete media for 30 minutes. Cells were fixed with 4% PFA and stained for VE-Cadherin (Santa Cruz Biotechnology, #6458, 1:200) with a secondary antibody coupled with Alexa-Fluor-488.

### Scratch wound assay

24h after siRNA transfection cells were re-plated into a scratch wound assay device (IBIDI). On the following day a cell free gap of 500μm was created by removing the insert of the device. Images were taken immediately after removing the insert (0h) and after 16h. The cell free area was measured in Fiji and used to calculate the percentage of wound closure at 16h.

### RNA extraction and quantitative real time-polymerase chain reaction

RNA was extracted using the RNeasy Mini Kit (Qiagen) according to the manufacturer’s instructions. For HUVECs transfected with adenoviruses carrying *YAP* and *TAZ* gain of function mutations, 2µg of total RNA were reverse transcribed to cDNA using M-MLV reverse transcriptase (Invitrogen). For HUVECs transfected with siRNAs 90ng of RNA were reverse transcribed using RevertAid First Strand cDNA Synthesis Kit (ThermoFisher Scientific). qRT-PCR was performed using TaqMan reagents and probes (Applied Biosystems) (listed in Supplementary Table 2). qRT-PCR reactions were run on a StepOnePlus real-time PCR instrument (ThermoFisher Scientific) or Quant Studio 6 Flex (Applied Biosystems) and expression levels were normalised to human *ACTB* or human *HPRT1* using the 2deltaCT method.

### Western blot

Protein was extracted from HUVECs using M-PER protein extraction reagent with Halt Protease and Phosphatase inhibitors (Pierce). Proteins concentration was assessed using a BCA protein assay kit (Pierce). Proteins were separated by SDS–PAGE and blotted onto nitrocellulose membranes (Bio-Rad). Membranes were probed with specific primary antibodies and then with peroxidase-conjugated secondary antibodies. The following antibodies were used: YAP 63.7 (Santa Cruz Biotechnology, sc-101199, 1:1000), GAPDH (Millipore, MAB374, 1:4000). The bands were visualized by chemiluminescence using an ECL detection kit (GE Heathcare) and a My ECL Imager (Thermo Scientific).

### Dual luciferase reporter assay

Renilla-luciferase reporter assays for TEF-1 (48), RBPj (49), BRE (50, 51) and FOPflash (52)-Luciferase promoter activity were performed as follows: 48 hours after gene knockdown by siRNA HUVECs were cotransfected with 600 ng of Luciferase reporter gene construct and 300 ng of pRL-CMV (Promega) using Lipofectamine2000 and incubated for 4 hours. Cell extracts were prepared 72 hours post siRNA transfection and 24 hours post Luciferase reporter transfection, and luciferase activity was measured using a dual luciferase system as described (53). Experiments were carried out in duplicates and results were normalized to the correspondent FOPflash/Renilla measurement.

### Microarray and gene set enrichment analysis

Microarray studies were performed as described(46). In brief, total RNA was extracted from HUVECs using the RNeasy kit (Qiagen) and RNA quality assessed with the 6000 nano kit and an Agilent Bioanalyser. RNA was labelled according to the Affymetrix Whole Transcript Sense Target Labeling protocol. Affymetrix GeneChip® Human Gene 2.0 ST arrays were hybridized and scanned using Affymetrix protocols. Data were analysed using the Affymetrix expression console using the RMA algorithm; statistical analysis was done using DNAStar Arraystar 11. Heat maps of gene signatures were plotted using RStudio, Inc.

### Statistical Analysis

Statistical analyses were performed using GraphPad Prism software and statistical significance was determined using unpaired Student *t*-test. When technically possible the investigators were blinded to genotype during experiments and quantification.

## Author Contribution

FN designed the study, performed experiments, analyzed the results and wrote the manuscript; AKB, YTO, ACV, AS and JRC performed experiments, analyzed the results and reviewed the manuscript; IH and EBK performed experiments; SA performed quantitative analysis and analyzed results; CAF provided critical feedback. HG and MP designed the study and wrote the manuscript.

## Acknowledgments

We thank members of the Vascular Biology (Berlin) and Vascular Patterning (VIB – Leuven) Laboratories for helpful discussions. We thank the Cancer Research UK - London Research Institute and the Max Delbrück Center for Molecular Medicine Animal Facilities for animal care and technical support. We thank Dr. Axel Behrens for kindly providing the *Taz*^*fl/fl*^ mice. We thank Dr. Walter Birchmeier and Dr. Daniel Besser for providing the Normalizer and BRE-luc reporter, Dr. Eric Sahai and Dr. Nic Tapon for providing the TEF1-reporter, and Dr. Michael Gotthardt and Dr. Michael Radke for access to the Flexcell® Tension System and technical assistance. We specially thank Dr. Veronique Gebala, Dr. Andre Rosa and Dr. Baptiste Coxam for helpful comments on the manuscript.

FN was financially supported by the Fundação para a Ciência e a Tecnologia (FCT), CRUK-CRICK and the MDC. ACV and AKB were supported by the DZHK (German Centre for Cardiovascular Research), AS was supported by the EMBO (European Molecular Biology Organization), JRC was supported by the FCT. CAF is supported by the FCT, EC-ERC Starting Grant, Portugal2020 program. MP is supported by the Max Planck Society, the ERC Starting Grant ANGIOMET, the Deutsche Forschungsgemeinschaft, the Excellence Cluster Cardiopulmonary System, the LOEWE grant Ub-Net, the DZHK, the Stiftung Charité and the EMBO Young Investigator Program. HG is supported by the DZHK and ERC Consolidator Grant Reshape 311719.

---

**Figure 1 - Figure supplement 1.**
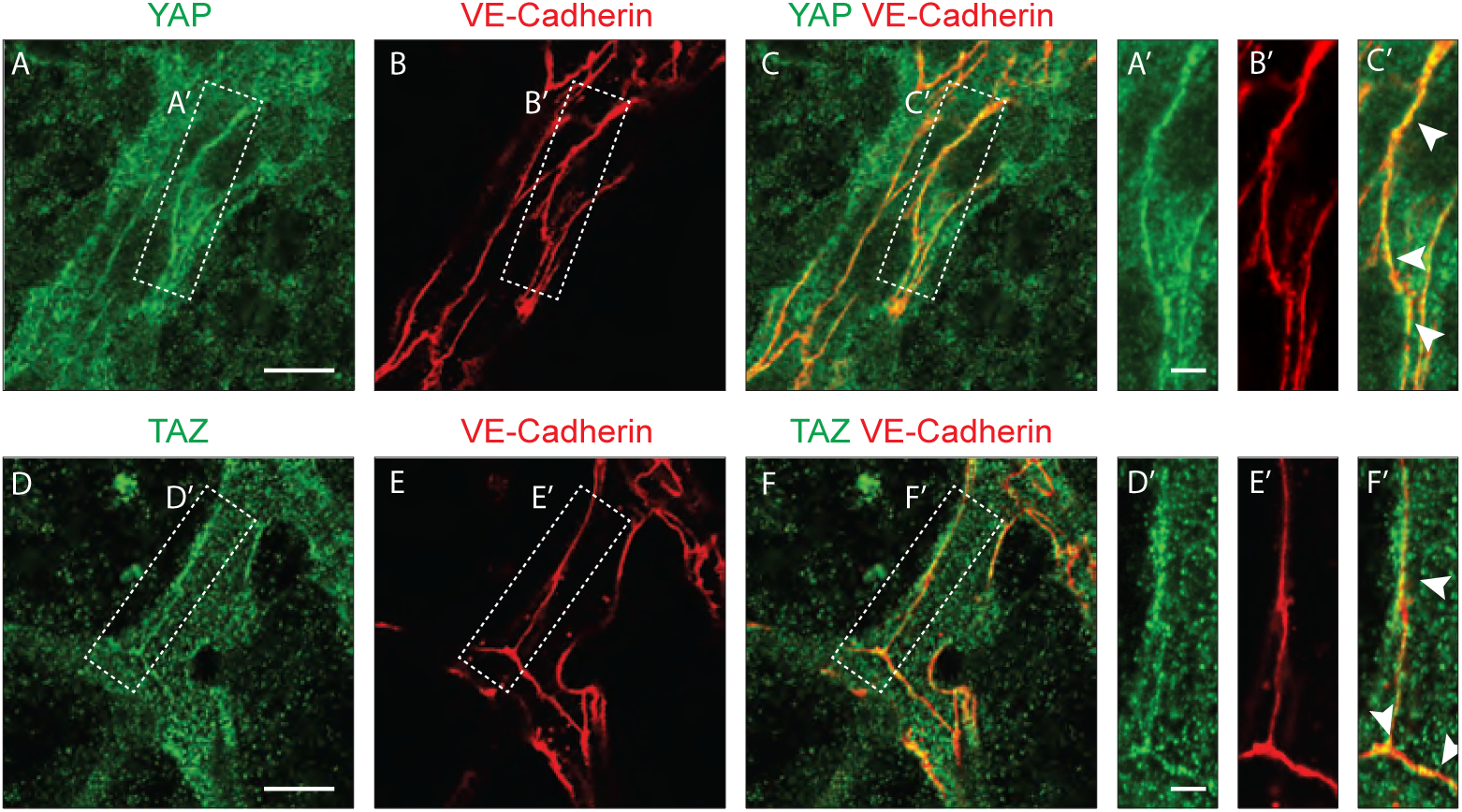
YAP and TAZ localise at endothelial adherens junctions in the mouse retina. Immunofluorescence stainings of YAP (green, A-C,A’-C’), TAZ (green, D-F, D’-F’) and VE-Cadherin (red, B,E,B’,E’) were performed in wild-type mouse retinas at P6. Arrows, co-localisation of YAP or TAZ with VE-Cadherin. Images correspond to single confocal planes. Scale bar A-C and D-F 10μm, A’-C’ and D’-F’ 3μm.

**Figure 2 - Figure supplement 1.**
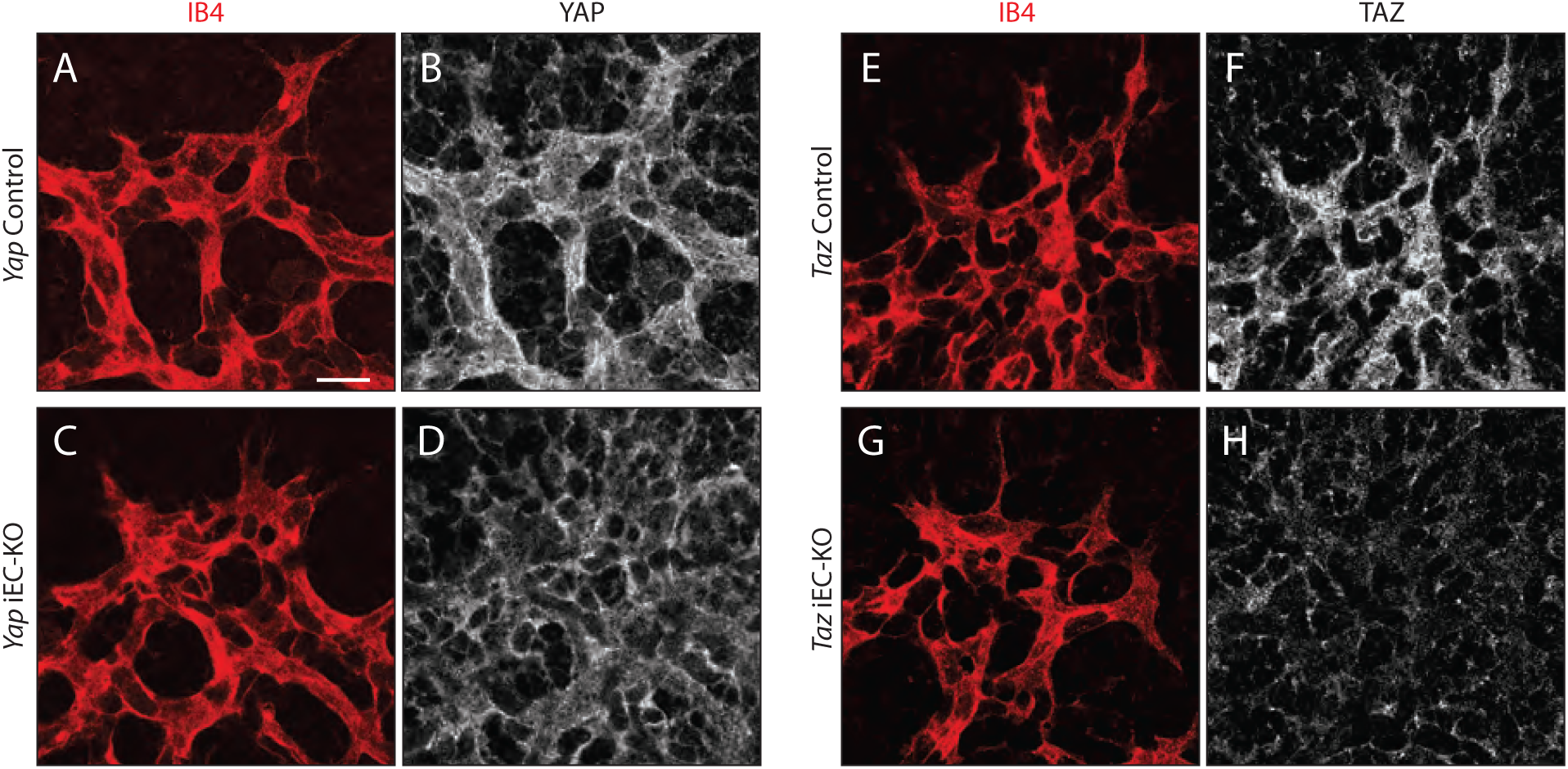
YAP and TAZ proteins are lost upon Cre-mediated genetic deletion in P6 mouse retinas. *Yap* iEC-KO (*Yap*^fl/fl^ Pdgfb-iCreERT2+/wt), *Taz* iEC-KO (*Taz*^fl/fl^ Pdgfb-iCreERT2+/wt) and respective littermate control mice (*Yap*Control, *Yap*^fl/fl^ and *Taz*Control, *Taz*^fl/fl^) were injected with tamoxifen at P1 and P3. At P6, mouse retinas were stained for YAP (grey, B,D), TAZ (grey, F,H), and with Isolectin B4 (IB4; red, A,C,E,G). Scale bar: 20μm.

**Figure 2 - Figure supplement 2.**
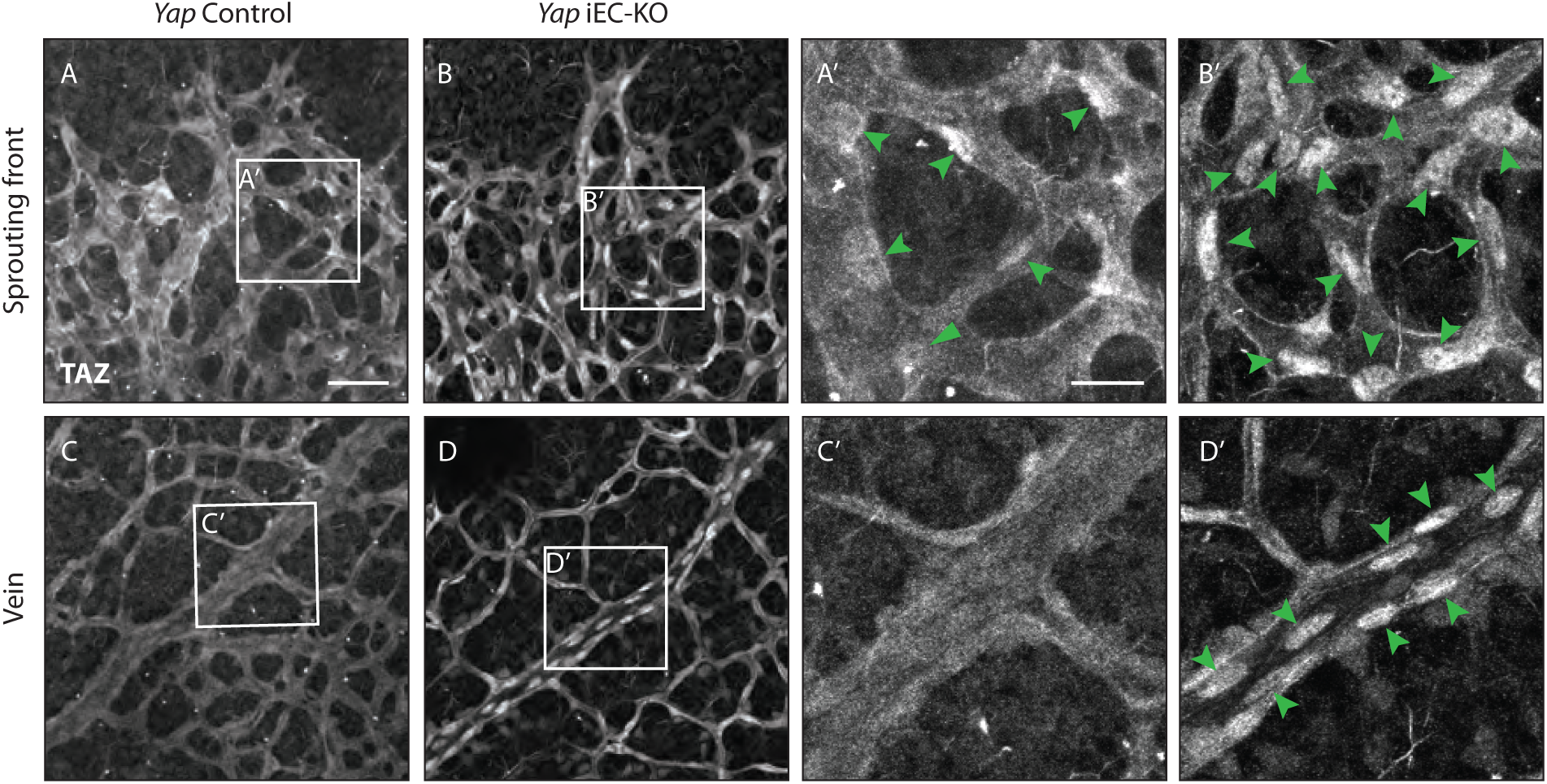
TAZ compensates for the loss of YAP in endothelial cells *in vivo*. Retinas from P6 *Yap* iEC-KO (B,D and B’,D’) and littermate controls (A,C and A’, C’) were immunostained for TAZ. Green arrowheads, nuclear Taz. A,B, A’,B’ images correspond to maximum projection of z stack. C,D,C‘,D’ correspond to single confocal planes. Scale bar: A-D 50μm, A’-D’ 20μm.

**Figure 3 - Figure supplement 1.**
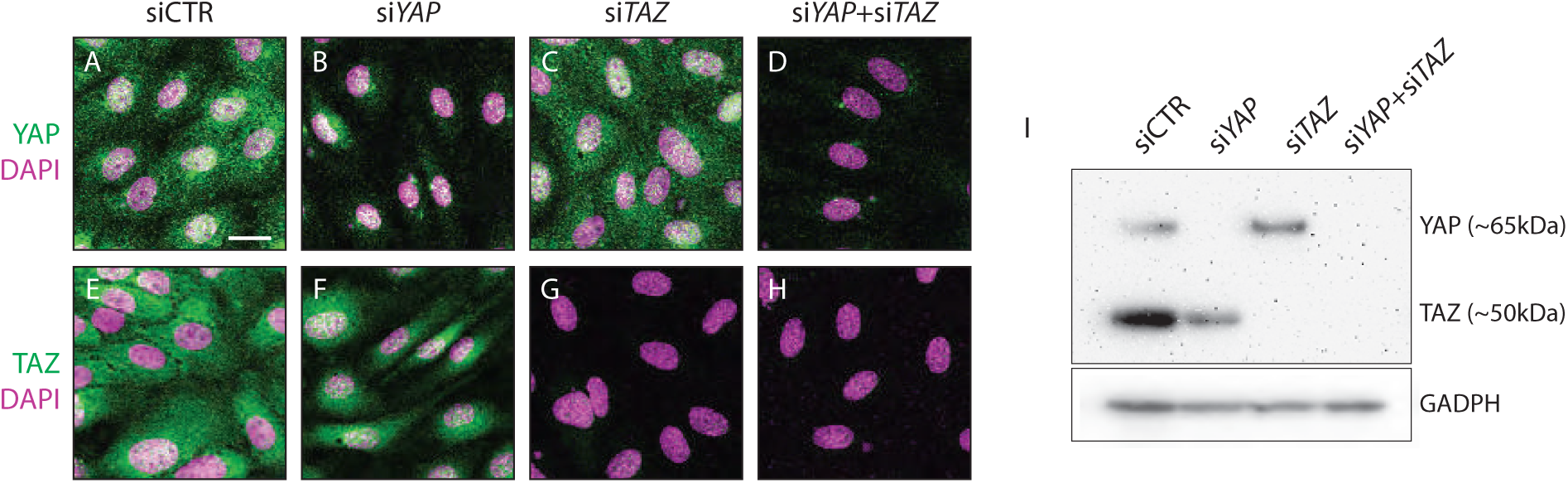
YAP and TAZ proteins are lost after gene knockdown by siRNA in HUVECs. HUVECs were treated with non targeting siRNA (siCTR) or siRNA targeting *YAP*, *TAZ* and *YAP* +*TAZ* for 24h. **A-H**, Immunofluorescence staining for YAP (green, **A-D**) or TAZ (green, **E-H**) and labelling of nuclei with DAPI (magenta) 72h after siRNA transfection. Scale bar: 10μm. **I**, Western blot for YAP/TAZ and GAPDH 72h after siRNA transfection.

**Figure 3 - Figure supplement 2.**
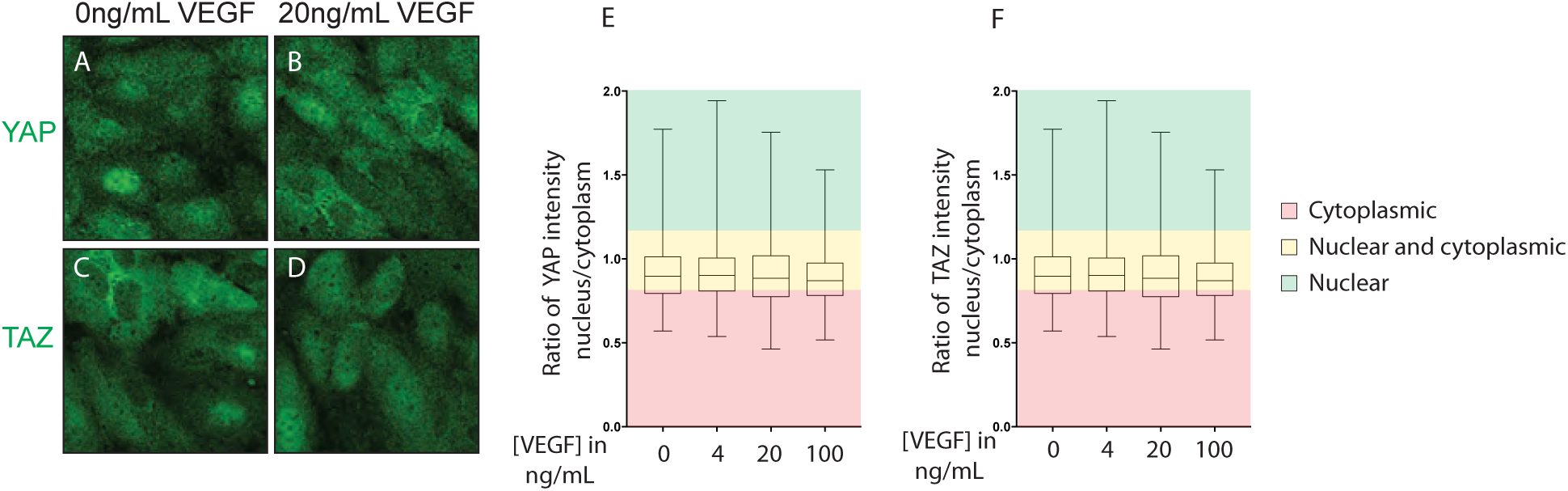
VEGF treatment does not affect YAP and TAZ subcellular localisation. Confluent HUVECs were treated with 0, 4, 20 or 100ng/mL of VEGF for 6h. **A-D**, Immunofluorescence staining for YAP (**A,B**) and TAZ (**C,D**) after 6h of 0 and 20 ng/mL of VEGF treatment. **E,F,** Quantification of YAP (**E**) and TAZ (**F**) subcellular localisation with increasing concentrations of VEGF treatment. Data represent ratio of YAP (E) or TAZ (F) staining intensity in the nucleus/cytoplasm. Cytoplasmic, ratio< 0.8; nuclear and cytoplasmic, ratio between 0.8-1.2; nuclear, ratio >1.2. Box and whiskers graph of one representative experiment out of 3. Whiskers are min to maximum. n= 3 independent experiments; > 500 cells analysed per condition per experiment.

**Figure 4 - Figure supplement 1.**
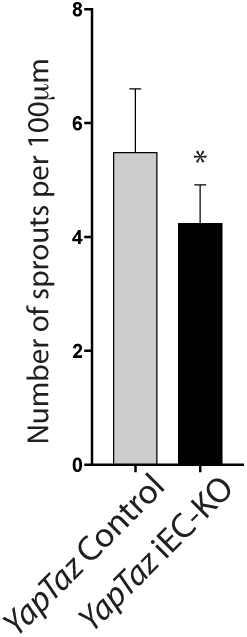
Combined loss of YAP and TAZ leads to decreased number of sprouts in the developing mouse retina. Quantification of number of sprouts per 100 μm of sprouting front extension at P6 in *YapTaz* iEC-KO (n= 9 pups) and littermate control mice (n=9 pups). Data are mean + SD. p values were calculated using unpaired t-test. *, p<0.05.

**Figure 7 - Figure supplement 1.**
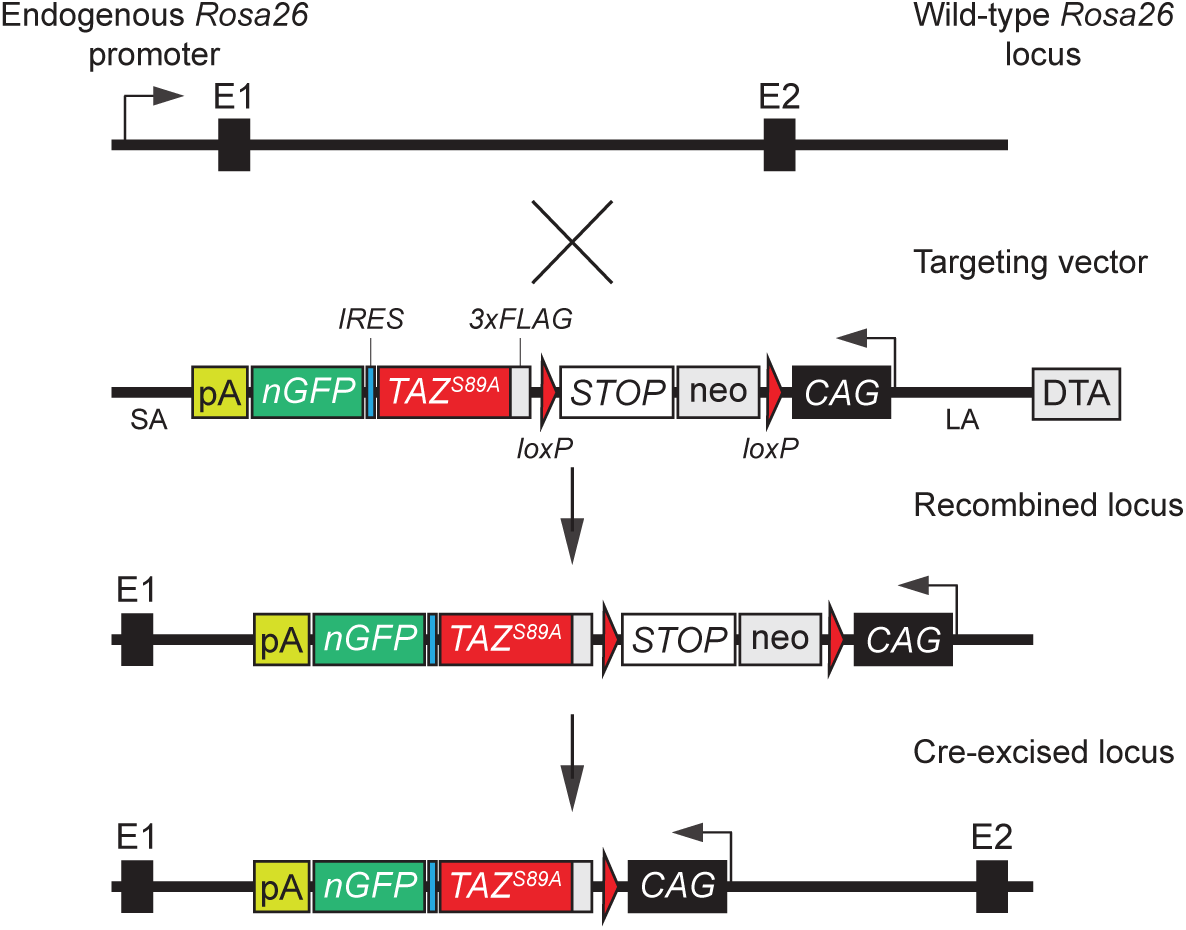
Targeting strategy used for the generation of the conditional TAZ gain-of-function mouse model. cDNA coding for a 3xFLAG-tagged human TAZ S89A was inserted into a Rosa26 targeting vector downstream of the ubiquitous CAG promoter. The cDNA also included an internal ribosome entry sequence (IRES) and a nuclear-localized enhanced green fluorescence protein (nEGFP) for monitoring transgene expression. To allow Cre-dependent expression of 3xFLAG-TAZS89A and of the EGFP reporter, a floxed transcriptional STOP cassette was incorporated between the 3xFLAG-TAZS89A-IRES-nEGFP sequence and the CAG promoter.

**Figure 7 - Figure supplement 2.**
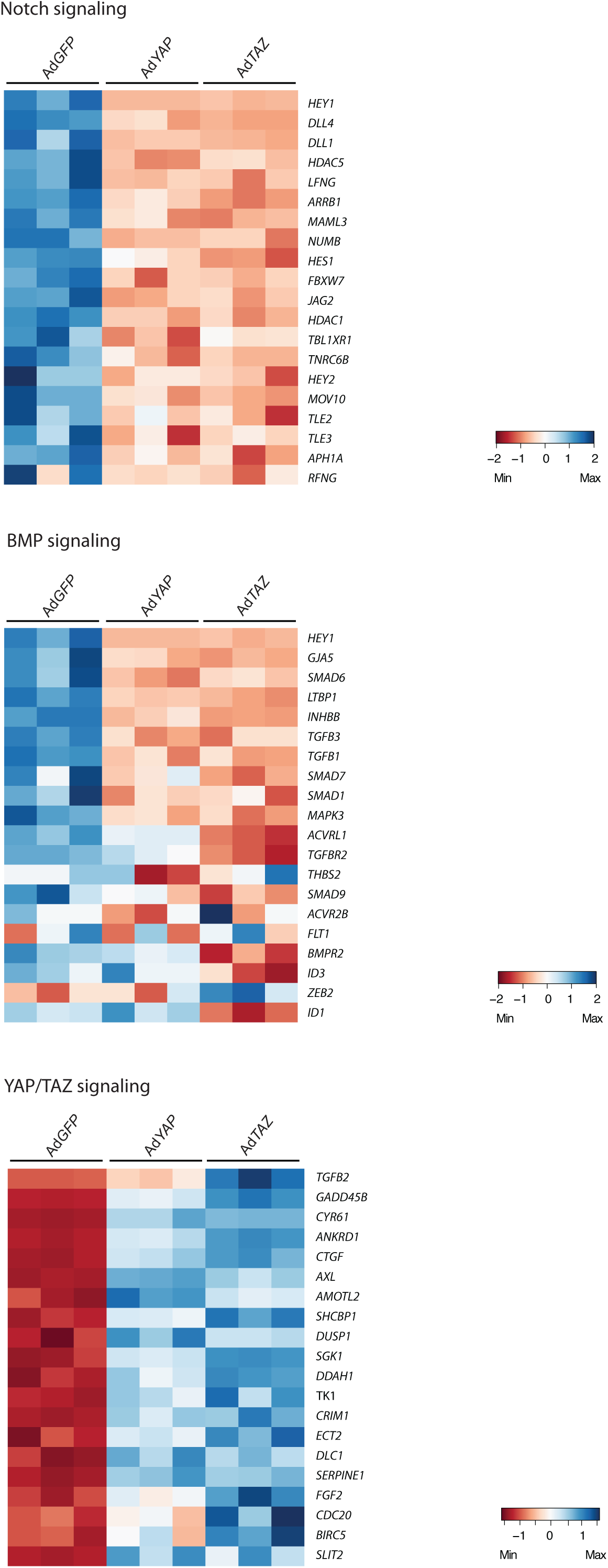
Microarray of YAP and TAZ gain of function mutant cells. Heatmaps of Notch, BMP and Hippo pathway genes in control (Ad*GFP*), Ad*YAP* and Ad*TAZ* HUVECs.

**Figure 7 - Figure supplement 3.**
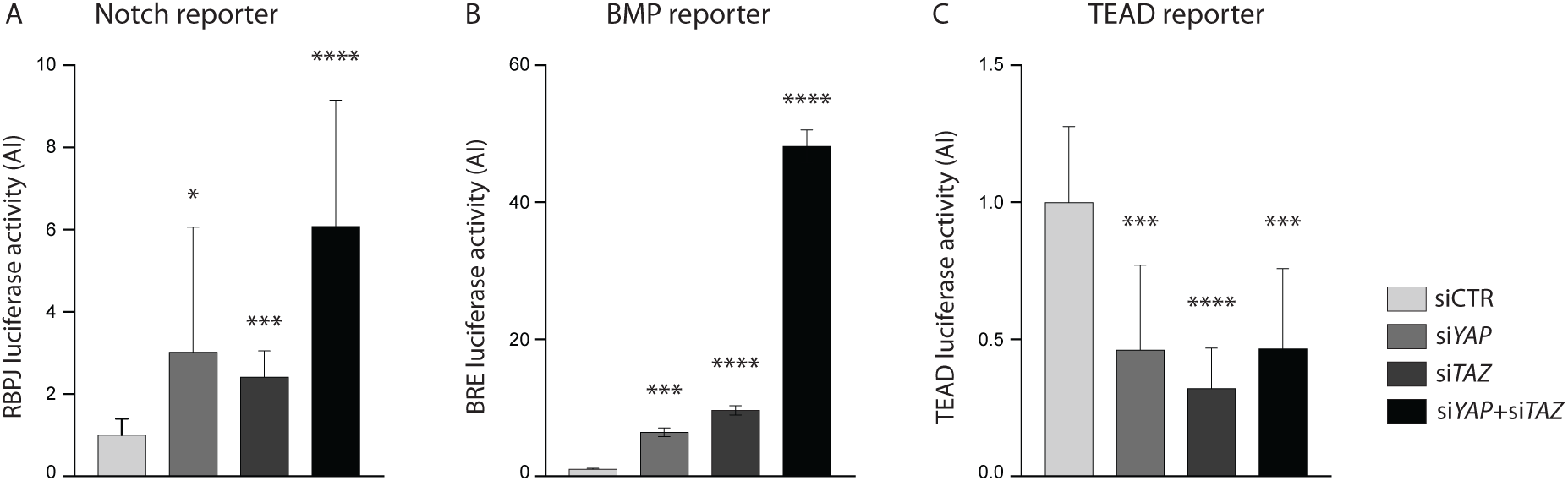
YAP and TAZ knockdown increases Notch and BMP reporter activities *in vitro*. **A-C**, Luciferase reporter assays in YAP, TAZ and YAP/TAZ knockdown HUVECs and controls for Notch reporter (A), BMP reporter (B) and TEAD reporter (C). Data are mean + SD. *p* values were calculated using unpaired *t*-test. n ≥ 3 experiments for Notch reporter, 3 experiments for BMP reporter, ≥ 6 experiments for TEAD reporter. *, *p*<0.05; **, *p*<0.01; ***, *p*<0.001; ****, *p*<0.0001.

**Figure 7 - Figure supplement 4.**
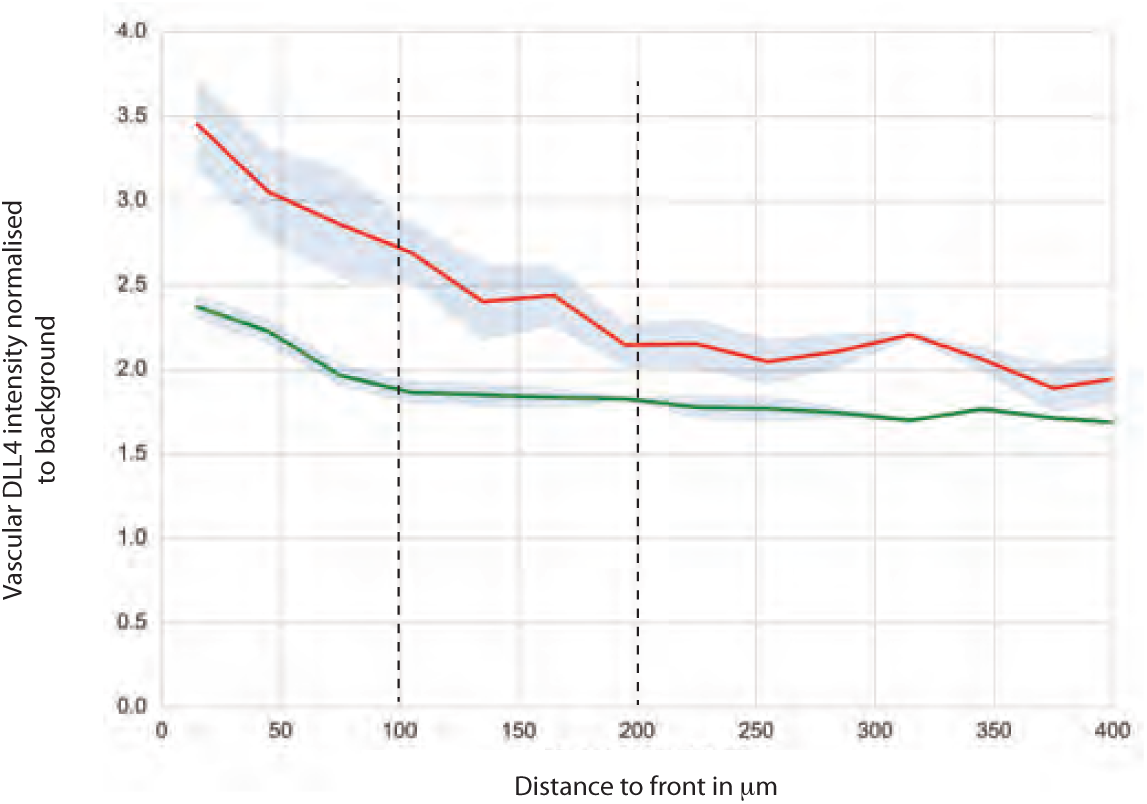
DLL4 intensity in *YapTaz* iEC-KO. Graph shows mean DLL4 staining intensity in the vascular retina of control (green) and *YapTaz* iEC-KO (red) P6 pups normalised to the background intensity. Data are mean +/- SEM. n= 3 control and 3 *YapTaz* iEC-KO.

**Supplementary table 1.**
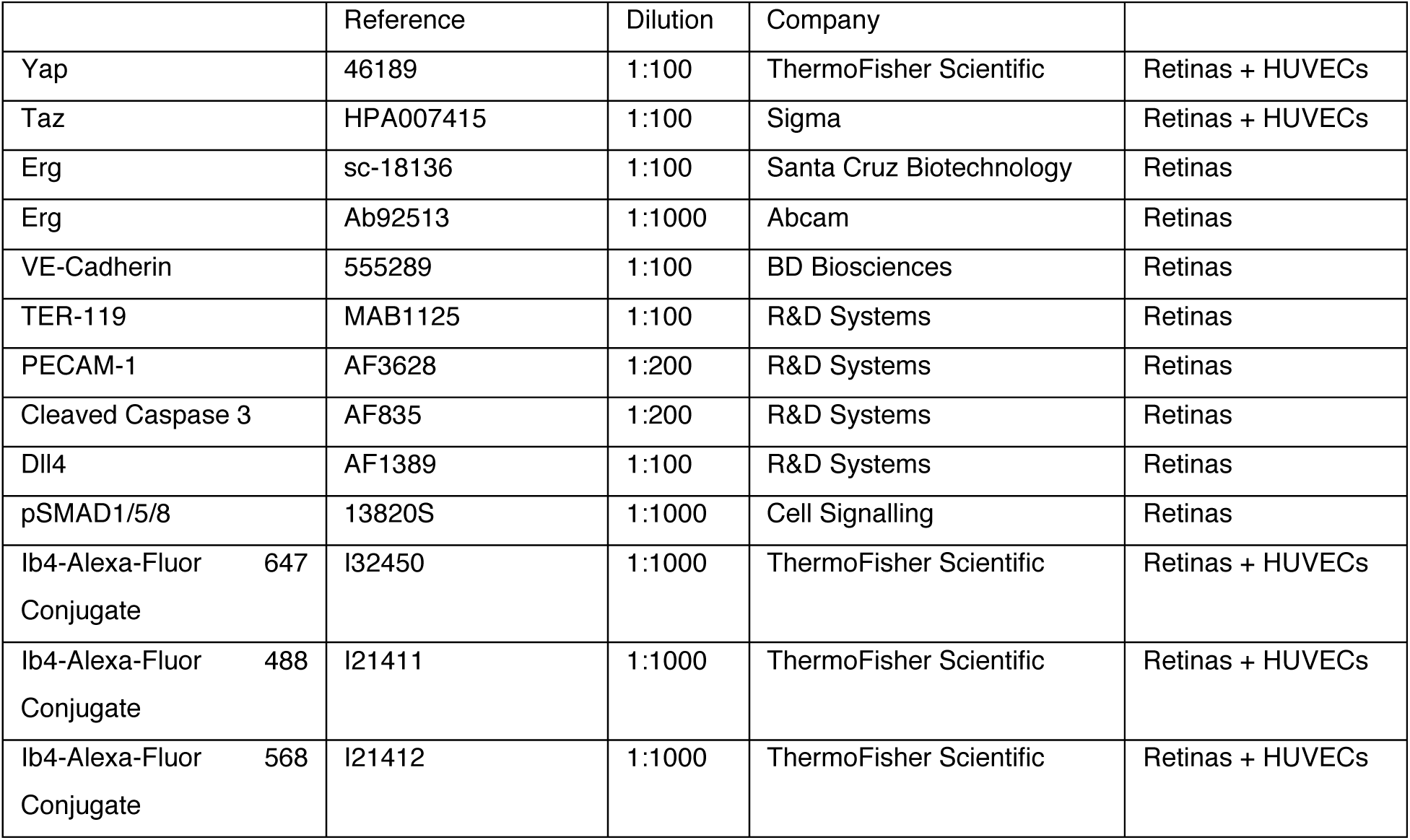
List of primary antibodies used.

**Supplementary table 2.** List of the TaqMan primers (Applied Biosystems) used.

| Target | Assay ID |
| --- | --- |
| LFNG | Hs00385436_g1 |
| DLL4 | Hs00184092_m1 |
| NRARP | Hs04183811_s1 |
| Hes1 | Hs00172878_m1 |
| Hey1 | Hs01114113_m1 |
| SMAD6 | Hs00178579_m1 |
| ENG | Hs00923996_m1 |
| UNC5B | Hs00900710_m1 |
| ID1 | Hs03676575_s1 |
| ID3 | Hs00171409_m1 |
| CTGF | Hs00170014_m1 |
| CYR61 | Hs00998500_g1 |
| ANKRD1 | Hs00923599_m1 |
| INHBA1 | Hs01081598_m1 |
| HPRT1 | Hs02800695_m1 |
| ACTB | Hs99999903_m1 |

